# Cell-level integration of gene expression and B cell receptor data captures complementary B cell information

**DOI:** 10.64898/2026.10.16.759689

**Authors:** Edel Aron, Gisela Gabernet, Burhan Sabuwala, Gur Yaari, Steven H. Kleinstein

## Abstract

Single-cell B cell analyses commonly define relationships among cells using gene expression (GEX), although paired B cell receptor (BCR) sequences provide complementary information about receptor features, differentiation, and clonal relationships. We evaluated multiple approaches for integrating paired GEX and BCR data into cell-level representations and compared their ability to capture complementary aspects of B cell biology. We compared BCR representations based on language-model embeddings and engineered sequence features across multiple integration strategies. Spaces dominated by BCR language-model embeddings captured broad sequence and clonal information but could distort cell type relationships, as assessed using independent cell surface protein measurements. In contrast, concatenating GEX with selected BCR features (isotype, somatic hypermutation frequency, and CDRH3 length) incorporated receptor information while retaining transcriptional and surface-protein-associated structure. The resulting space reduced discordance between GEX-based cell type annotations and BCR characteristics and revealed B-cell substructure supported by both transcriptional and receptor-derived features. Thus, integrating GEX and BCR data captures complementary biological information and provides a richer foundation for single-cell B cell analysis.

## Introduction

Single-cell RNA sequencing (scRNA-seq) enables transcriptome-wide profiling of individual cells and has transformed the characterization of cellular types and states. However, gene expression (GEX) captures only part of a cell’s biological state, motivating multi-omics approaches that measure complementary features in the same cells [1]. For example, Cellular Indexing of Transcriptomes and Epitopes by Sequencing (CITE-seq) profiles cell-surface proteins using antibody-derived tags (ADTs) alongside GEX [2]. Because ADTs directly measure surface proteins, they can provide clearer and more interpretable markers for cell type annotation than the corresponding transcripts [3]. Yet other modalities can provide complementary information not captured by either transcripts or protein markers. In B cells, paired B cell receptor (BCR) sequencing provides information regarding receptor features, differentiation history, and clonal relationships that can be linked to the transcriptional state of the same cell.

Single-cell profiling has enabled increasingly detailed characterization of immune-cell heterogeneity [4–8]. B cells are central to adaptive immunity, differentiating from naïve cells into memory B cells and antibody-secreting cells (ASCs) following antigen encounter. Each B cell expresses a B cell receptor composed of an immunoglobulin heavy chain (IGH) paired with either a kappa (IGK) or lambda (IGL) light chain. The variable regions contain V, D (heavy chain only), and J gene segments that form the antigen-binding site, whereas the heavy-chain constant region determines isotype. Platforms such as 10x Genomics can simultaneously profile GEX and paired BCR sequences from individual cells [9].

BCR sequences encode information that is not directly available from GEX, including isotype, somatic hypermutation (SHM) frequency, clonal relationships, and the length and physicochemical features of the heavy chain complementarity-determining region 3 (CDRH3) [10]. Isotype and SHM frequency are particularly informative for B cell subtyping; naïve B cells typically co-express IgD and IgM and generally lack SHM, whereas class switching and the accumulation of SHM generally accompany antigen-driven differentiation. Other BCR features also vary among B cell populations. For example, naïve B cells tend to have longer CDRH3 regions than differentiated cells, although their distributions overlap [11–13]. These features are not independent: isotypes differ in their CDRH3 characteristics, mutation frequencies vary among V genes, and CDR3 length is associated with gene usage [14–16].

Transcript, protein, and BCR measurements capture different aspects of B cell identity and do not always support the same cell type assignment. For example, some memory B cells lack *CD27*, a marker commonly used to distinguish memory from naïve B cells [17]. Joint analysis of GEX and adaptive immune receptor repertoire data can reveal immune cell populations and dynamics that are difficult to resolve using either modality alone [5, 18–21]. Similarly, integrating GEX and ADT measurements can improve cell type annotation [22], but ADT measurements are limited to the selected antibody panel and do not capture many BCR-specific features.

BCR data can therefore complement GEX-based annotation and support repertoire-informed classification [13]. However, BCR information alone is also insufficient to define cellular state; clonally related cells can occupy distinct transcriptional states, while cells from different populations may share similar receptor features [18]. Thus, neither GEX nor BCR alone provides a complete representation of B cell state, motivating cell-level approaches that integrate information from both modalities.

Several methods, including Benisse [23], BiGCN [24], CoMBCR [25], and Ibex [26], jointly analyze BCR and GEX data for applications such as cell type annotation, clustering, and developmental inference. Existing approaches differ in their objectives and representations, but commonly emphasize CDR3 sequences, heavy chains, or clone-level relationships. It remains unclear which cell-level representations most effectively integrate transcriptional and receptor-derived information while preserving biologically meaningful organization.

Here, we evaluated multiple approaches for integrating paired GEX and BCR data into cell-level representations and compared their ability to capture complementary aspects of B cell biology. We show that integrating GEX with selected BCR features introduces complementary receptor information while retaining transcriptional and surface-protein-associated structure, reduces discordance between GEX-based annotations and BCR features, and resolves B cell substructure that is less apparent from GEX alone.

## Results

### GEX-based distances are not sufficient for B cell clustering and type annotation

GEX-based distances are commonly used to cluster cells. To determine whether such distances could benefit from integrating BCR data, we investigated the quality of B cell type annotations that they can produce. Starting with scRNA-seq data from 16 individuals profiled in a study of type 1 diabetes [27], Leiden clustering of B cells identified 11 clusters, which were then annotated using CellTypist [28] with cluster-level majority voting (Figure 1A-B). This analysis identified 91,021 naïve B cells, 19,664 memory B cells, and 1,028 antibody-secreting cells (Figure 1C-D). However, paired BCR data revealed that ∼7% of the cells annotated as naïve were potentially misannotated as they expressed BCRs inconsistent with a naïve cell type, i.e., their BCRs were class-switched or somatically hypermutated. This was not due to the number of clusters, as such misannotations were frequent across a range of clustering resolutions (Supplementary Figure 1). Thus, GEX-based distances are not sufficient for cluster-based identification of a major B cell type.

**Figure 1.**
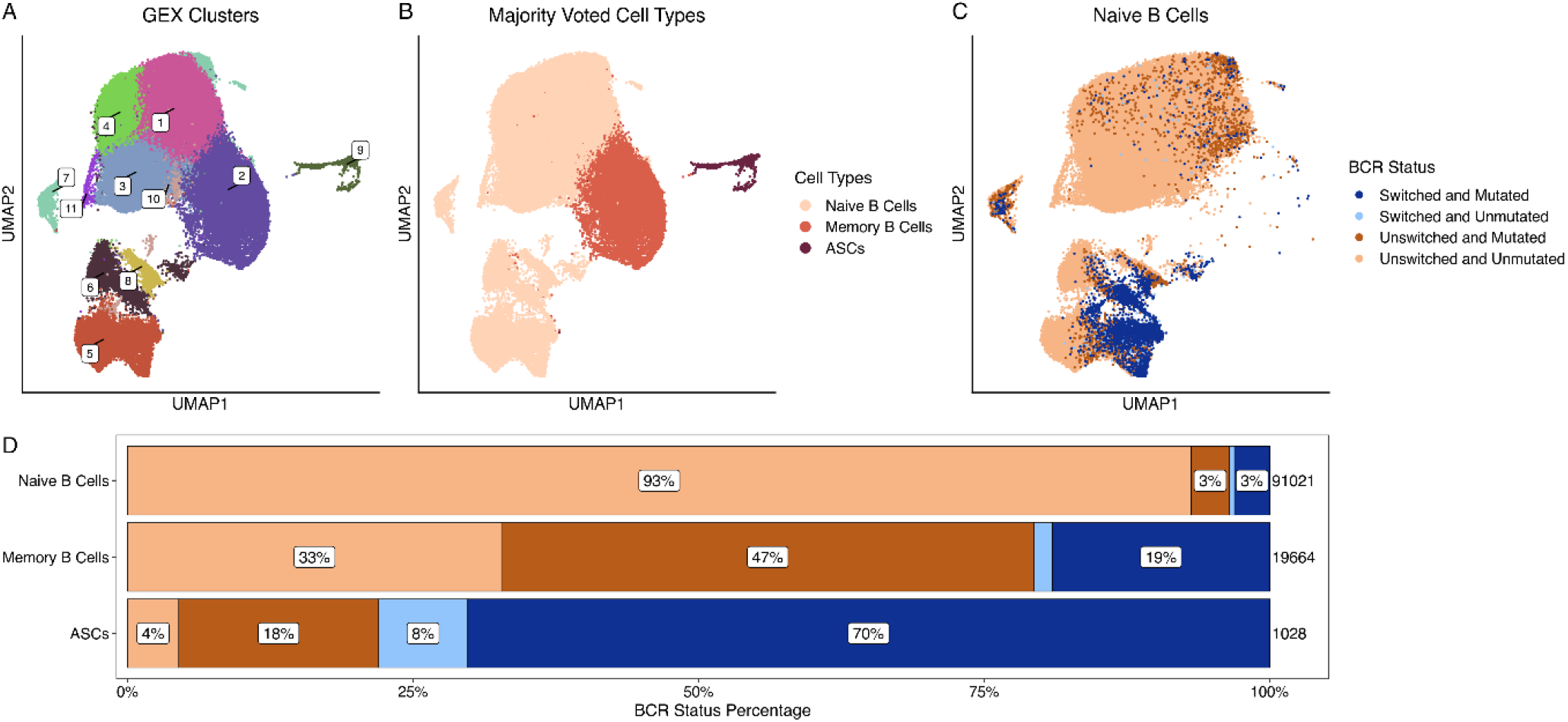
BCR features are inconsistent with cluster-based B cell annotations. **A)** UMAP of the GEX space, colored by the 11 clusters identified by Leiden clustering (resolution 0.4). **B)** The same UMAP colored by cell types, as annotated by CellTypist with cluster majority voting and the Adult Human Blood reference. **C)** The same UMAP restricted to naïve B cells and colored by BCR status. **D)** Stacked bar plot showing the composition of each cell type by BCR status (colored the same as C). Total cell counts for each cell type are provided to the right of each bar.

To evaluate GEX-based distances independently of the clustering method, we used CellTypist to annotate individual cells (Supplementary Figure 2A), then analyzed their 20 nearest neighbors (hereafter referred to as the “local neighborhood”) for consistency. Our dataset contained 88,277 naïve B cells, 22,670 memory B cells, and 766 ASCs (Supplementary Figure 2B-C). Among the naïve B cells, ∼97% expressed BCRs that were consistent with their naïve annotation. The local neighborhoods of these cells were also consistent with their annotation. 100% of the neighbors (median of 20/20 neighbors) were labeled as naïve by CellTypist and all (median of 20/20 neighbors) expressed BCRs with features that were consistent with being a naïve B cell (Supplementary Figure 2D-E). In contrast, there was a lack of consistency in the local neighborhoods of the 3% (2,521 of 88,277) of B cells annotated as naïve by CellTypist that expressed class-switched and/or mutated BCRs. These cells expressed BCRs with features inconsistent with a naïve annotation (somatic hypermutation and/or class switched isotypes), yet only 20% of their neighbors (median of 4/20 neighbors) were labeled as non-naïve by CellTypist and 40% (median of 8/20 neighbors) expressed BCRs with features that suggested a non-naïve status (Supplementary Figure 2D-E). These patterns were stable across a range of neighborhood sizes (Supplementary Figure 3). Thus, neither cluster-level annotation nor local neighborhoods defined by GEX-based distances adequately resolved the discordance between cell type annotations and BCR characteristics, motivating the integration of both modalities.

### Language-model embeddings reffect multiple BCR sequence features

To facilitate the integration of GEX and BCR information, we converted BCR sequences into fixed-length numerical representations. As one approach, we represented each BCR as a vector of three features already known to be associated with B cell type: isotype, SHM frequency and CDRH3 length [29–31]. Language-model embeddings offer an alternative approach. These embeddings simultaneously reflect multiple BCR features such as IGHV usage and efficiently transfer to downstream tasks such as specificity prediction [32]. We evaluated five BCR-specific or protein language models that generate fixed-length embeddings from amino acid sequences: AntiBERTa2 [33], AntiBERTy [34], BALM-paired [35], ESM2 [36], and Immune2vec [37] (Table 1). All of the embedding methods organized cells according to five sequence-derived features that we evaluated, albeit to different degrees: IGHV gene subgroup (a set of genes sharing at least 75% nucleotide identity [38]), light chain locus, isotype, SHM frequency, and CDRH3 length (Figure 2; Supplementary Figure 4). This organization was expected because these features are directly encoded in, or closely associated with, the BCR sequence. While some of these features, such as isotype and SHM frequency, were also apparent in the GEX-based embeddings, others such as IGHV gene subgroup and CDRH3 length, were only seen in the language-model embeddings. Thus, language-model embedding of the BCR can provide information beyond the cell’s GEX profile. We selected the concatenation of three features (isotype, SHM frequency and CDRH3 length), and a representative embedding method (AntiBERTa2) for downstream integration analyses.

**Figure 2.**
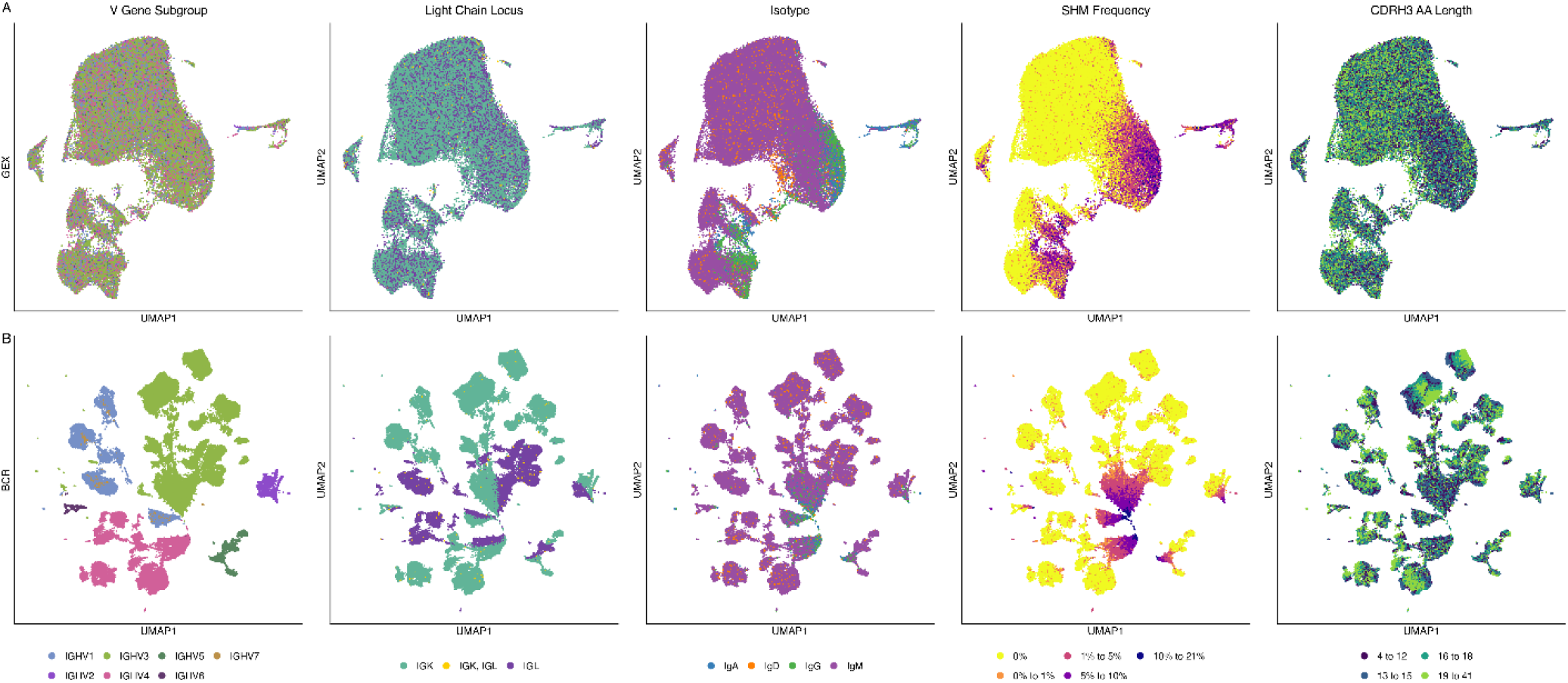
BCR embeddings preserve key BCR features. UMAPs of GEX **(A)** and BCR (AntiBERTa2) spaces **(B)**. Each column is colored by a BCR feature, from left to right: V gene subgroup, light chain locus, isotype, binned SHM frequency, and binned CDRH3 amino acid length.

**Table 1.** Embedding models. Asterisks denote the best performers according to neighbor matching.

| Model | Based On | Dimensions | Max Seq Length | Chains trained on | Understands Pairing? | Groups By |
| --- | --- | --- | --- | --- | --- | --- |
| AntiBERTa2 | RoFormer | 1024 | 256 | H, L, H+L (separately) | No | Light chain locus*, SHM frequency*, V gene subgroup* |
| AntiBERTy | BERT | 512 | 510 | H, L, H+L (separately) | No | Light chain locus, V gene subgroup* |
| BALM-paired | RoBERTa | 1024 | 510 | HL, LH (concatenated) | Yes | Isotype*, light chain locus, SHM frequency* |
| ESM2 (650M parameter) | BERT | 1280 | 512 | H, L, H+L, HL/LH (concatenated) | No | Light chain locus*, V gene subgroup* |
| Immune2vec | Word2Vec | 100 (configurable) | - | H, L, H+L + (warning) HL/LH (concatenated) | No | SHM frequency*, V gene subgroup |

### Integrated spaces improve distances between cells with similar BCR features

We defined three spaces as controls to compare our integration approaches to: (1) GEX alone (Supplementary Figure 2A), (2) GEX without filtering of IG-related genes (referred to as GEX+IG) (Figure 3A), and (3) BCR alone, using the AntiBERTa2 embedding (Figure 3B). Then, we evaluated three approaches for integrating GEX and BCR data: (1) an adaptation of Seurat’s weighted nearest neighbors (WNN) framework using GEX and BCR k-nearest-neighbor graphs (Figure 3C), (2) concatenation of GEX principal components and BCR-embedding principal components (Figure 3D) referred to as “concatenation of embeddings” (CE), and (3) concatenation of GEX principal components with BCR features (Figure 3E) referred to as “concatenation of features” (CF). For CE and CF, each modality was rescaled to contribute equal total variance, so that neither would dominate because of its dimensionality or scale.

**Figure 3.**
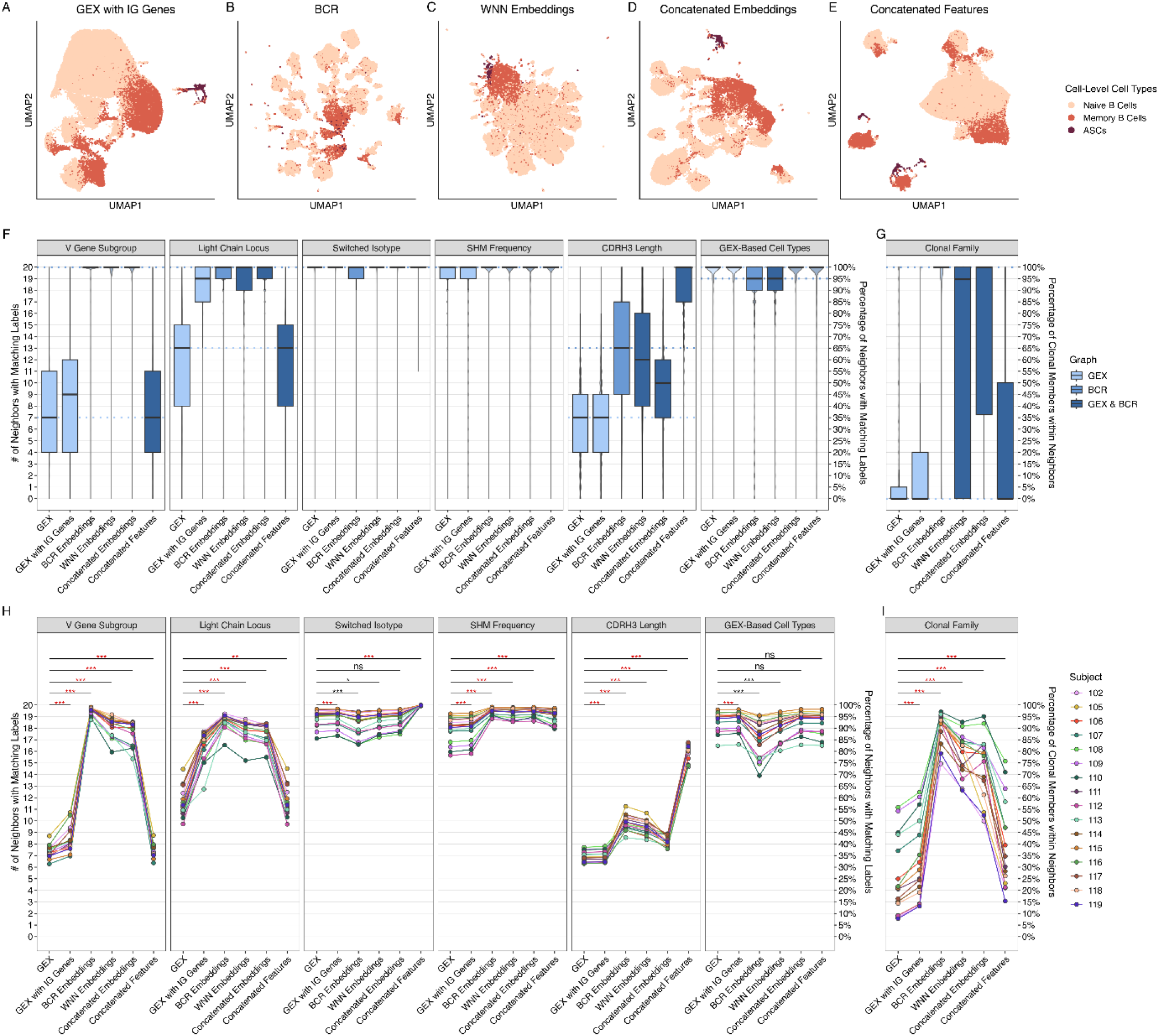
Integrated spaces preserve both BCR and GEX features within local neighborhoods. **A-E)** UMAPs based upon five spaces, colored by cell-level cell type annotations: **A)** GEX with IG genes retained (i.e., IG-related genes were not removed prior to finding the most variable features, running PCA and calculating neighbors), **B)** BCR embeddings (AntiBERTa2), **C)** WNN embeddings, **D)** concatenated embeddings, and **E)** concatenated features. **F)** Box plots showing the number (left axis) and percentage (right axis) of each cell’s k = 20 nearest neighbors that share the same BCR feature (V gene subgroup, light chain locus, isotype, SHM frequency, CDRH3 amino acid length within 2 amino acids) or GEX-based cell type annotation, as indicated on each panel, for each space. Violin plots behind each box plot show the full distribution of values. Boxes span the interquartile range (IQR) with the median line; whiskers extend to 1.5x IQR. **G)** Box plot calculated as in (F), but for clonal relationships among cells (for expanded clones only). Percentages are relative to the smaller of the clone size minus one and k. **H-I)** Per-subject means of the values in (F) and (G), respectively. Lines connect values from the same subject across spaces. Each space was compared with GEX using a paired Wilcoxon signed-rank test across subjects (***P < 0.001; **P < 0.01; *P < 0.05). Red and black asterisks indicate significant increases or decreases relative to GEX, respectively.

We evaluated each cell’s local neighborhood in each space for consistency of BCR features (Supplementary Figure 5). First, we considered five BCR features as discrete values: IGHV gene subgroup, light chain locus (IGK and/or IGL), isotype (class-switched vs. unswitched), SHM status (mutated vs. unmutated), and CDRH3 length. We also considered clonal relationships (subject-specific clone IDs that identify members of the same clonal family). As expected, GEX alone generally showed the least consistency in the local neighborhood, with significantly fewer neighbors sharing the same IGHV gene or CDRH3 length compared with other spaces (Figure 3F; Supplementary Figure 6). While light chain locus and SHM status were better captured by GEX alone, with more than half of neighbors sharing the same feature, this was still significantly lower than in virtually all other spaces. Retaining the IG genes in the GEX data generally improved the local neighborhood consistency for all features. BCR alone, WNN and CE had similar performance, with moderate to high local neighborhood consistency across all BCR features (Figure 3F,H). Clonal relationships followed a similar pattern (Figure 3G,I); among cells in expanded clones, a median of 100% of possible clonal neighbors were clonally related in the BCR and CE spaces (with a median of 95% in WNN), as compared with 0% in GEX alone. In contrast to the embeddings-based spaces, CF strongly retained the three BCR features explicitly included in its construction (isotype, SHM frequency, and CDRH3 length) and was more similar to GEX alone for other features that were not explicitly encoded in the feature vector. Despite clonal identity not being encoded in its features, the CF space also significantly increased the fraction of clonally related neighbors relative to GEX, although its median remained at 0%. Thus, integrating GEX with BCR data generally improved the local neighborhood consistency of BCR features.

We next evaluated CDRH3 length, SHM frequency and CDRH3 physicochemical features (measured by 5 Atchley and 10 Kidera factors) in a continuous manner by calculating the Spearman correlation between each cell’s value and the mean value among its nearest neighbors. In all of the BCR-containing spaces (BCR, WNN, CE and CF), CDRH3 length correlations were substantially higher in all BCR-containing spaces (0.748-0.990) compared with GEX alone (0.312) or GEX+IG (0.315) (Figure 4A). SHM frequency correlations showed more modest increases in the BCR-only and concatenated spaces (0.713-0.726 versus 0.675 for GEX and 0.680 for GEX+IG), while WNN was slightly lower than GEX (0.663; Figure 4B). BCR-containing spaces also showed increased local correlations for Atchley or Kidera factors (Supplementary Figure 7). Together, these analyses show that including BCR information in an integrated space tends to bring cells with similar BCR features closer together.

**Figure 4.**
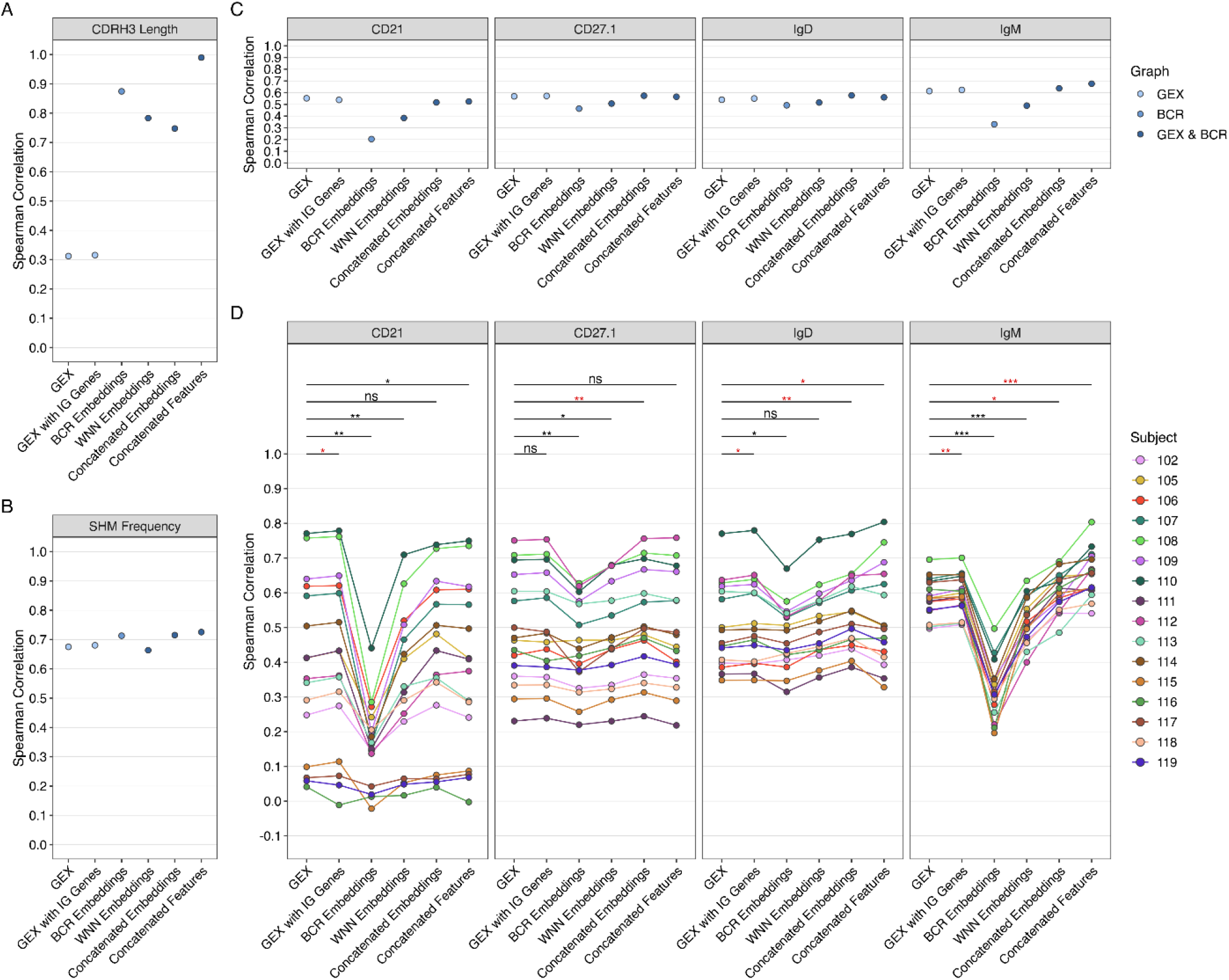
Local neighborhood consistency of BCR features and independent ADT measurements across spaces. **A-B)** Spearman correlation between each cell’s (A) CDRH3 amino acid length or (B) SHM frequency and the mean value among its 20 nearest neighbors, for each of the six spaces. **C)** Spearman correlation between each cell’s expression of four B-cell ADT markers (CD21, CD27.1, IgD, IgM) and the mean expression among its k = 20 nearest neighbors, calculated across all cells. **D)** Per-subject Spearman correlations for the four ADT markers in C across the six spaces. Lines connect values from the same subject. Each space was compared to GEX using a paired Wilcoxon signed-rank test across subjects (***P < 0.001; **P < 0.01; *P < 0.05). Red and black asterisks indicate significant increases or decreases relative to GEX, respectively.

### Spaces based on concatenation of BCR information do not compromise cell type relationships

Members of the same clonal family can span multiple cell types, such as memory B cells and ASCs [39, 40]. Thus, spaces driven too strongly by BCR sequence features or clonal relationships could obscure biologically meaningful cell type differences. We investigated whether the increased local neighborhood consistency achieved with integrated spaces came at the expense of a compromised GEX-associated cellular structure.

We first evaluated local neighborhoods for cell type consistency. Cell type annotations were based on CellTypist run on individual cells without clustering. Using GEX alone, 100% of local neighbors (median of 20/20 neighbors) were labeled as the same cell type (Figure 3F). As expected, the local neighborhood consistency of spaces based on BCR alone was significantly worse, although a median of 19/20 neighbors were still annotated with the same cell type (Figure 3F, H). The integrated spaces CE and CF had 100% of local neighbors (median of 20/20 neighbors) annotated with the same cell type, while 95% of local neighbors (median of 19/20 neighbors) were the same under the WNN space (Figure 3F). There was no significant difference between GEX alone and the CE or CF integrated approaches (Figure 3H). Interestingly, GEX+IG showed a small, but significant, increase in the local neighborhood consistency of cell type annotations compared with GEX alone. Overall, these results suggest that integrated spaces retain broad cell type relationships.

As the cell type annotations from CellTypist were derived from GEX, we sought a way to evaluate how well each space reflected cell type structure independently of GEX. To accomplish this, we leveraged single-cell measurements of surface protein expression from antibody-derived tags (ADT). We focused on four cell surface markers with known associations with B cell types: CD21, CD27, IgD and IgM [17]. For each of these markers, we calculated the Spearman correlation between protein expression in each cell and the mean expression among its nearest neighbors (Figure 4C; Supplementary Figure 8). Under this framework, betterspaces are expected to have higher correlations, indicating that cells with similar surface-protein expression are nearer to one another, and thus that similar cell types are closer together. We observed qualitatively similar results when using other metrics instead of Spearman correlation such as Pearson correlation, although the magnitudes of the scores varied by metric (Supplementary Figure 9).

Consistent with the observation that many different cell types can express BCRs with the same or highly similar amino acid sequences (e.g., members of a clonal family), the BCR alone space showed significantly lower cell surface protein correlations compared with GEX across all four protein markers (0.20-0.49 versus 0.54-0.61). WNN also underperformed GEX for CD21, CD27, and IgM, likely because its automatically learned weights favored the BCR modality (90,296 cells favored the BCR modality, 21,395 cells favored the GEX modality, and 22 cells were tied). This result illustrates that adding BCR information does not necessarily produce a space well suited to cell type analysis if the integration is dominated by sequence-derived structure. In contrast, the cell surface protein neighborhood correlations for CE and CF were comparable to (and for some markers significantly higher than) those obtained using GEX alone or GEX+IG (Figure 4D).

As isotype can be inferred directly from the BCR sequence, it was noteworthy that the BCR alone space had significantly lower neighborhood correlations for both IgD and IgM protein measurements when compared to GEX alone or GEX+IG. This is likely because the embeddings we evaluated only encoded the V(D)J region and not the constant region which defines the isotype. Including the constant region as part of the embedding caused the neighborhood correlations for all embeddings-based spaces to increase for IgD and to significantly increase for IgM (Supplementary Figure 10). We note that these correlations were still lower than GEX alone for the BCR and WNN spaces.

Finally, we tested how sensitive these results were to parameter choices for the integration approaches (i.e. the relative weighting of the two modalities, the concatenation strategy and the number of principal components), assessed using both local neighborhood consistency and cell surface protein correlations. For the concatenation approaches, varying the relative weight of GEX from 0 (BCR only) to 1 (GEX only) changed CE and CF mainly at the extremes; results were very stable over a range of weights around equal weighting (Supplementary Figure 11). We also explored three approaches for concatenation: combining raw embeddings or features with GEX prior to dimensionality reduction (“raw”), combining raw embeddings or features with GEX PCs (“reduced_gex”, used for CF), and combining BCR PCs with GEX PCs (“reduced_both”, used for CE). None of the alternative concatenation strategies improved the integrated spaces (Supplementary Figure 12). Compared to its alternatives, the CE approach we used throughout this study had the highest local neighborhood consistency for all BCR features except CDRH3 length, as well as higher cell surface protein correlations. Similarly, the CF approach used throughout this study had similar or slightly higher local neighborhood consistency and cell surface protein correlations compared to its alternatives (except for CDRH3 length). Finally, we found that local neighborhood consistency and cell surface protein correlations in the CE and CF spaces were relatively stable once more than five principal components from each modality were used (Supplementary Figure 13).

Overall, these results indicate that spaces based on concatenation of BCR information (CE and CF) can achieve increased local neighborhood consistency of BCR features without compromising cell type relationships. CF was selected for subsequent analyses because it retained both cell type structure and valuable BCR features well while allowing the contribution of individual BCR features to be interpreted directly.

### Integrated GEX and BCR features improve annotation consistency and resolve B cell substructure

Having shown that the CF space increases the local neighborhood consistency of BCR features, we sought to determine whether this space would also improve cell type annotation. We previously found ∼7% of B cells (6,271 of 91,021) annotated as naïve by CellTypist with cluster-level majority voting in the GEX space expressed BCRs that were inconsistent with this label (i.e., they were class switched or contained SHM) (Figure 1D).

When cells were annotated using CellTypist at the cell level instead, we found that 2,521 cells annotated as naïve B cells expressed inconsistent BCR features. Analysis of the local neighborhoods of these cells suggested that using the CF space for cluster-based cell type annotation could reduce this discordance. In the CF space, the local neighborhood of these cells included more cells annotated by CellTypist as non-naïve compared to the GEX alone space (median 10 vs. 4 neighbors, Figure 5A). It also included more cells with non-naïve BCR features (median 20 vs. 8 neighbors, Figure 5B). In fact, all integrated spaces separated unswitched and unmutated naïve-labeled cells from switched and/or mutated cells more effectively than GEX alone (Supplementary Figure 14). The CF integrated space thus organized these cells into neighborhoods enriched for other cells with features that should produce a more accurate cell type annotation.

**Figure 5.**
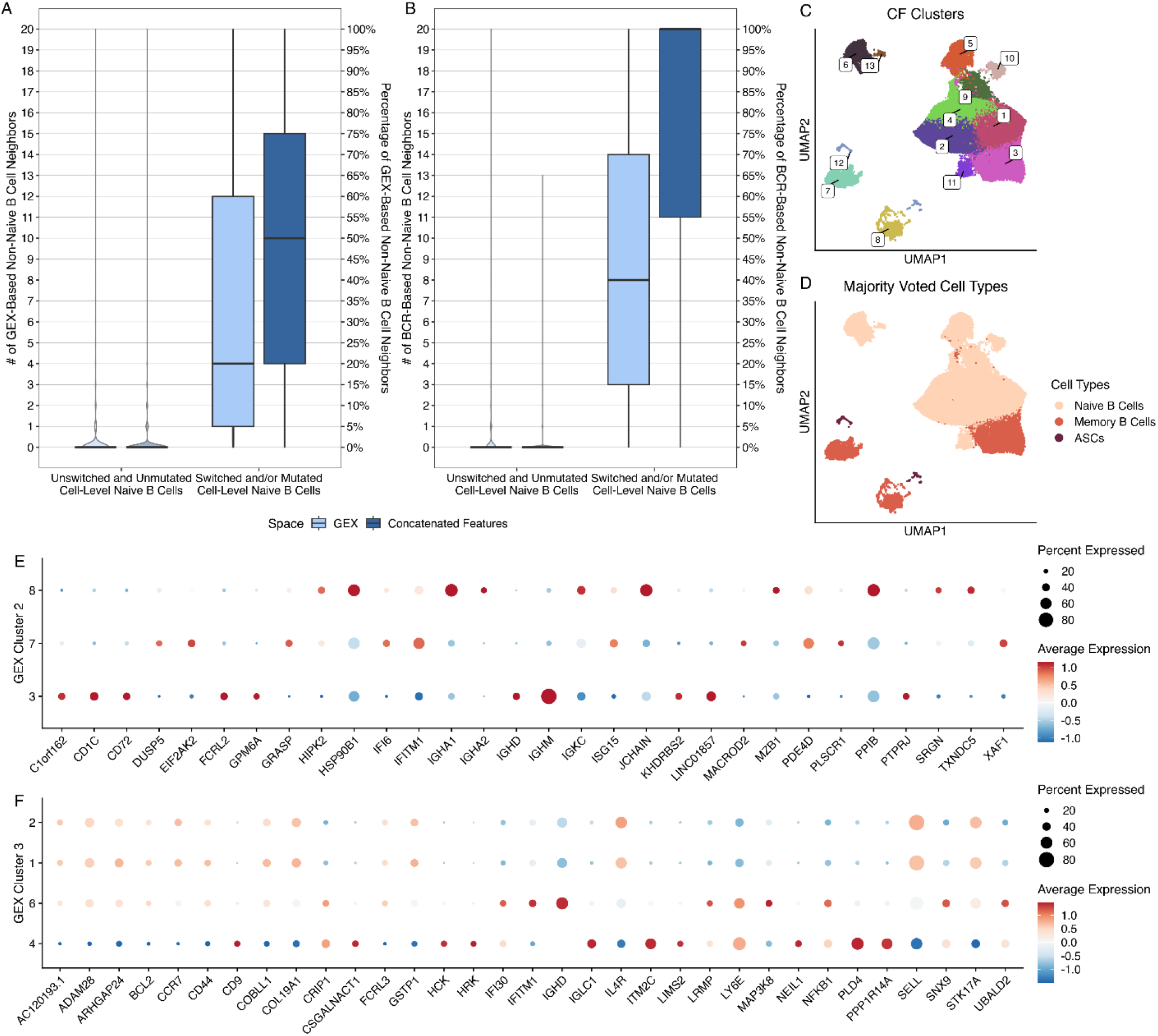
Cell type annotation in the CF space reduces discordance between BCR characteristics and cell-type labels. **A)** Number (left axis) and percentage (right axis) of each naïve B cell’s k = 20 nearest neighbors that were labelled as non-naïve B cells by CellTypist at the cell-level in the GEX (light blue) and CF (dark blue) spaces. Naïve B cells are grouped by BCR status as unswitched and unmutated (left), or switched and/or mutated (right). **B)** Same as in (A), but for neighbors with non-naïve BCR characteristics (switched and/or mutated). **C)** UMAP of the CF space, colored by the 13 clusters identified with Leiden clustering (resolution 0.4). **D)** The same UMAP colored by cell type, assigned by CellTypist with cluster majority voting. **E)** Dot plot of the top differentially expressed genes among CF clusters 3, 7, and 8, which received most cells from GEX cluster 2. **F)** Dot plot of the top differentially expressed genes among clusters 1, 2, 4, and 6, which received most cells from GEX cluster 3. In (A) and (B), boxes span the interquartile range (IQR) with the median line; whiskers extend to 1.5x IQR. In (E) and (F), dot size indicates the percentage of cells expressing each gene, and color indicates scaled average expression.

We next tested whether cluster-level annotation in the integrated space reduced inconsistency between cell type annotations and BCR features. Leiden clustering in the CF space yielded 13 clusters, which we annotated using CellTypist with cluster-level majority voting (Figure 5C-D). Not only did this CF-based annotation increase the total number of naïve B cells (from 88,277 at the cell-level to 90,374 cells), but fewer of these cells expressed BCRs with features that were inconsistent with this annotation. None of the B cells annotated as naïve in the CF space expressed a class-switched BCR, compared to 3.5% of naïve B cells annotated in the GEX space. Overall, only ∼3% of the B cells annotated as naïve in the CF space expressed BCRs inconsistent with this label (compared to ∼7% when clustering in the GEX space). In addition, consistent with their inactivated status, fewer of these naïve B cells were members of expanded clones (2.37% vs. 2.68% for the GEX space). Together, these findings suggest that working in the CF space reduces discordances in naïve B cell annotations.

Memory B cells were also distributed across three clusters in the CF space, corresponding predominantly to IgM^+^ (cluster 3), IgG^+^ (cluster 7), and IgA^+^ (cluster 8) memory cells. This separation likely reflected the fact that isotype was explicitly included as a feature in the CF space. Nevertheless, differential gene expression analysis comparing these three clusters revealed transcriptional differences beyond isotype (Figure 5E). Cells in the IgA^+^ memory cell cluster expressed higher levels of JCHAIN, MZB1, and TXNDC5, consistent with increased secretory pathway activity. Cells in the IgG^+^ memory cell cluster showed increased expression of interferon-regulated genes, including IFI6, IFITM1, and ISG15 [41]. Finally, cells in the IgM^+^ memory cell cluster expressed higher levels of CD72 and FCRL2, in addition to IGHM, consistent with memory B cells [42]. They also expressed CD1C, a marker typically associated with marginal zone B cells. Thus, analysis in the CF space identified memory cell subsets that reflected both isotype and transcriptional differences.

To further investigate whether clusters in the CF space better reflected biologically relevant differences, we annotated the individual cells using the Allen Institute for Immunology’s PBMC B cell reference [43]. This analysis found that the CF space could concentrate cell subtypes such as transitional B cells. For example, the cluster with the highest proportion of transitional B cells in the GEX space was cluster 3 (59%; Supplementary Figure 15). Virtually all of the cells in this cluster (99.6%) were divided between four clusters in the CF space (clusters 1, 2, 4 and 6), with cluster 4 having a much higher proportion of transitional B cells (73%). Differential gene expression analysis between these four clusters revealed that cluster 4 (which contained 62.5% of the cells from GEX cluster 3) also showed increased expression of transitional B cell markers, including CD9, ITM2C, PLD4, and PPP1R14A (Figure 5F). Clusters 1, 2 and 6 also had distinct gene expression profiles. Cluster 6 showed differential expression of 72 genes, including increased expression of several interferon-regulated genes (IFI30, IFITM1, LY6E), as well as IGHD, NFKB1, and SNX9. While clusters 1 and 2 were less transcriptionally distinct (both upregulated CCR7 and SELL), they showed differential expression of 10 genes and 6 genes, respectively (Figure 5F). Together, these observations suggest that the CF space separates transitional B cells from neighboring naïve-like populations that were intermixed in the GEX space.

## Discussion

Single-cell gene expression (GEX) and B cell receptor (BCR) sequencing capture complementary aspects of B cell biology. GEX reflects cell type and activation state, whereas BCR sequences encode receptor specificity, differentiation history, and clonal relationships. Here, we explored approaches for integrating these modalities into a common cell-level representation that more effectively captures B cell biology. We found that a joint space integrating GEX with selected BCR features reduced discordance between GEX-based cell type annotations and BCR features and revealed B cell population substructure that was less apparent from GEX alone.

Single cell GEX profiles form the basis for many analyses, including cell type annotation. However, some cell types are hard to distinguish based on transcriptional profiles alone. Reanalyzing data from patients with type 1 diabetes along with healthy controls, we found that many cells annotated as naïve based on the GEX profiles expressed BCRs with features that were inconsistent with this cell type. This observation of misannotated cells was also found in a second dataset comprised of individuals with systemic lupus erythematosus and healthy controls (Supplementary Figure 16). The integration of GEX and BCR data to create a joint space for analysis offers the possibility for improved cell type annotation. While the specific problem of identifying naïve B cells could be partially addressed by simply removing the naïve label from cells with mutated and/or class switched BCRs, it is unclear what label to then apply to these cells. Focusing on discrete labels may also miss important B cell population substructure, which can have overlapping states or form a continuous trajectory. An integrated space offers the possibility of capturing these differentiation trajectories using pseudotime or related developmental trajectory approaches [44], and may be especially beneficial if transcriptional and BCR changes capture processes occurring on different timescales [25].

The distance between cells in an integrated space is expected to reflect similarity in both transcriptional profiles as well as BCR features. Indeed, we found that the local neighborhood surrounding each cell in integrated spaces tended to express BCRs with the same isotype, mutation status and CDRH3 length. However, the specific set of features reflected in the integrated space was highly dependent on the BCR representation used. When using engineered features (CF space), only the specific features included in the representation influenced the space. Thus, for example, the CF space brought together clonally related cells far less than the embeddings-based spaces. In contrast, language-model embeddings retained broad sequence-derived features, including IGHV gene subgroup, light chain locus, and clonal relationships. Such features do not necessarily provide helpful information regarding a cell’s type or state and may even hinder cell organization in a biologically meaningful way. Clonally related cells can occupy distinct functional subsets (e.g., memory and ASC), as can cells expressing BCRs with the same IGHV gene.

To evaluate whether integrated spaces were organizing cells in a biologically meaningful way, we leveraged cell surface protein expression. Protein marker expression is the standard in immunology for defining cell types. We reasoned that cells with similar cell surface protein expression levels should be close together in a biologically meaningful space. This approach showed that concatenating GEX data with three engineered BCR features (isotype, somatic hypermutation frequency, and CDRH3 length) offered a favorable balance between maintaining this cell surface protein-associated organization while introducing BCR features related to differentiation. Unlike concatenating GEX data with language-model embeddings (CE space), this space also makes the factors influencing cell relationships explicit, which makes the space more interpretable. While some clusters in the CF space had strong BCR isotype associations, they also exhibited other transcriptional differences suggesting a broader biological relevance. Thus, the integrated space did more than reproduce its input BCR features and provided a basis for identifying biologically supported substructure.

Beyond cell type annotation, integrated spaces could support developmental or pseudotime analyses by combining transcriptional changes with directional BCR processes such as class switching and SHM accumulation. Prior studies have identified correspondence between B cell clonal lineages and transcriptionally inferred developmental trajectories [45, 46]. While the BCR embedding methods explored here tended to have a negative impact on cell type organization, future language-model representations could be adapted to emphasize task-relevant BCR features. Protein language models have been adapted to specialized problems such as enzyme representation and mutation-effect prediction as well as immune-repertoire analysis [47–49]. Fine tuning BCR models to prioritize cell type and state-associated features over gene usage or amino acid sequence similarity may improve their utility for multimodal integration. Light-chain information also warrants further investigation because the BCR and WNN spaces showed prominent organization by kappa and lambda chains, which have known physicochemical differences [50].

This study has several limitations. While we found consistent results in two human PBMC datasets generated using 10x Genomics, broader validation is needed across tissues, diseases, ages, and experimental platforms. BCR repertoires and B cell population structures vary with these factors [11, 29, 40, 43]; and GEX-based inference of immunoglobulin isotype may also depend on the reference genome and ancestry [51]. Although we tested a variety of integration parameter options and combinations, we did not exhaustively optimize them. Some of our results may be dependent on specific parameter settings and neighborhood-based analyses may be sensitive to the number of cells or technical factors like sequencing depth [52]. The cell filtering strategy may also have introduced biases. To compare embedding methods, we retained only cells successfully processed by all five methods, although analyses using a single embedding method or engineered features could retain more cells. Requiring paired heavy and light chain sequences excluded B cells affected by technical dropout, whereas filtering cells that were not annotated as B cells according to their GEX profile may have removed unusual B cell populations. Finally, the choice of which engineered features to be included in the integrated space will impact performance and may warrant further exploration. Additional BCR features, such as productive vs. sterile transcripts from methods such as sciCSR [53], could be highly informative. While the experiments analyzed here included targeted amplification and sequencing of the BCR, it is possible to apply this approach using BCR reconstructed from tools such as BraCeR or TRUST4 [54, 55].

In summary, we find that integrating GEX with BCR features creates a cell-level representation that captures complementary biological information unavailable from either modality alone. This joint GEX and BCR space reduces discordance between cell type annotations and BCR characteristics and offers a richer foundation for studying B cell identity, heterogeneity, differentiation, and receptor-associated biology.

## Methods

### Data collection and sample processing

Data from two studies were obtained from the NIH Gene Expression Omnibus (GEO) [56]. Both studies used 10x Genomics technology that included paired single-cell gene expression (GEX), antibody-derived tag (ADT), and B cell receptor (BCR) data. Sample collection and experimental procedures are described in the original studies [27, 57]. The primary dataset, hereafter referred to as Diabetes Nicholas, included 16 participants: five non-diabetic first-degree relatives, six autoantibody-positive individuals at risk of developing diabetes, and five individuals with recent-onset type 1 diabetes [27]. Peripheral blood mononuclear cell B cells were enriched using CD3^-^CD19^+^ sorting and antigen tetramers. The original study annotated cells as naïve, early memory, age- or autoimmune-associated, classical memory B cells, or plasmablasts, with the latter category potentially including plasma cells. Unless otherwise specified, the Methods describe analyses of the Diabetes Nicholas dataset. Processing of the second dataset, referred to as Lupus Siu, is described in the Supplementary Methods. A schematic overview of the analytical workflow is provided in Supplementary Figure 17.

### GEX data processing

For the GEX and ADT data, we used the filtered feature-barcode matrices generated by cellranger count that were deposited on GEO. The GEX count matrices were processed in R using Seurat v5.3.1 [58]. Sample-level matrices were merged into a single Seurat object containing 142,423 cells. Sample identifiers were appended to cell barcodes to ensure uniqueness. Cells expressing fewer than 200 genes or with mitochondrial transcript content greater than 20% were removed, leaving 142,173 cells. Counts were log-normalized using NormalizeData with a scale factor of 10,000. The 2,000 most variable genes were identified using the variance-stabilizing transformation method implemented in FindVariableFeatures and scaled using ScaleData.

Immunoglobulin (IG) and T cell receptor (TR) genes were excluded from the variable features because their inclusion can dominate downstream analyses and produce less biologically informative clusters [59]. Gene annotations were obtained from Ensembl v114 [60]. After removing IG and TR genes, 1,679 variable features remained. For comparison, we also generated a GEX space in which IG genes were retained but TR genes were excluded, leaving 1,894 variable features.

Principal component analysis (PCA) was performed using RunPCA with 50 components. Based on the inflection point of the elbow plot, the first 20 principal components (PCs) were used to construct approximate k-nearest-neighbor (kNN) and shared-nearest-neighbor graphs using FindNeighbors, with Euclidean distance and k = 20. The same PCs were used to generate uniform manifold approximation and projection (UMAP) visualizations with RunUMAP. Default parameters were used unless otherwise specified. Potential subject-specific batch effects were evaluated by inspecting UMAPs colored by subject. Batch correction was not applied to the Diabetes Nicholas dataset.

For cell type annotation, CellTypist v1.7.1, a logistic regression-based annotation tool [28], was applied using the Adult_Human_Blood model. For the main analyses, CellTypist labels were consolidated into three broad categories: naïve B cells; memory B cells, comprising age-associated, switched-memory, and unswitched-memory B cells; and antibody-secreting cells (ASCs), comprising plasmablasts and plasma cells. Sensitivity analyses used the CellTypist Immune_All_Low model, the Allen Institute for Immunology PBMC B cell reference [61], and Azimuth reference mapping [62] (Supplementary Figure 15).

Doublets were identified independently within each sample using scDblFinder with default parameters [63]. After doublet removal, 129,915 cells remained. Neighbor graphs and UMAP coordinates were recalculated on the filtered dataset.

### ADT data processing

ADT counts were stored as a separate assay in the Seurat object and normalized across cells using the centered log-ratio transformation. Only markers relevant to B cell biology were included in the evaluation; CD21, CD27, IgD, and IgM. Hashtag oligonucleotide and antigen-tetramer measurements were excluded.

### BCR data processing

For the BCR data, we used the AIRR rearrangement files (airr_rearrangement.tsv) that were deposited on GEO. BCR sequences were processed using nf-core/airrflow v4.3.1 [64], which incorporates tools from the Immcantation framework [65, 66]. V(D)J germline gene assignments were performed using IgBLAST v1.20.0 and the IMGT reference database bundled with nf-core/airrflow downloaded on November 26^th^, 2024 [67, 68].

We retained productive rearrangements with valid V and J gene assignments and junction lengths divisible by three. Cells containing only light chains, multiple heavy chains, unassigned heavy-chain constant regions, or ambiguous light chain assignments labeled as “multi” by Cell Ranger were excluded. D gene assignments were masked because they cannot generally be inferred reliably from short-read BCR sequences [69]. The Diabetes Nicholas dataset initially contained 242,088 BCR sequences. After processing with airrflow, 113,593 heavy-chain sequences were available for matching to GEX profiles. nf-core/airrflow identified clonal families separately within each subject. Sequences were first partitioned by V gene assignment, J gene assignment, and junction length. Junction nucleotide sequences were then hierarchically clustered, and clonal families were defined using a length-normalized Hamming distance threshold of 0.1.

SHM frequency was calculated within nf-core/airrflow as the proportion of nucleotide mismatches between each aligned heavy-chain V(D)J sequence and the inferred germline sequence. D gene segments were masked and not counted as part of the SHM frequency since D gene assignments can be unreliable in short-read data. Sequences with SHM frequency <1% were classified as unmutated and those with SHM frequency ≥1% as mutated, allowing for sequencing errors and undetected germline alleles absent from the reference database. CDRH3 physicochemical features were calculated from heavy-chain sequences using the aminoAcidProperties function in Alakazam v1.4.2. V, D, and J gene assignments and subgroup designations were extracted using the Alakazam getSegment function. Potential novel V gene alleles that could produce apparent mutations through reference misassignment were evaluated using TIgGER findNovelAlleles; no novel alleles were detected [70, 71].

### BCR sequence embeddings

BCR sequences were embedded separately for each sample using AMULETY v1.1 [75] and four publicly available models: AntiBERTa2 [76], AntiBERTy [34], BALM-paired [35], and ESM2 [36]. Heavy and light chains were included when supported by the model, as detailed in Table 1. AMULETY selected the light chain with the highest duplicate count when multiple light chains were associated with one heavy chain.

The effect of including the constant region was evaluated separately using AMULETY v2.1.2 and nf-core/airrflow v5.0.0 (Supplementary Figure 10). Immune2vec [37] was run independently using the recommended 100-dimensional representation and was trained on the analyzed dataset rather than on an external sequence corpus.

For each embedding model, sample-level embeddings were combined into a single cell-by-dimension matrix and matched to the Seurat object using cell barcodes. Embeddings that could not be matched to GEX profiles were excluded. To ensure direct comparability among methods, cells successfully processed by Immune2vec but not by all other embedding methods were also removed. This affected cells for which Immune2vec accommodated multiple light chains or a heavy chain without a paired light chain.

Each embedding matrix was stored as a separate BCR assay in a corresponding Seurat object. Because the embedding values did not require Seurat normalization or scaling, the original values were stored in both the data and scale.data layers to support downstream Seurat functions. PCA was applied to each BCR embedding matrix using 50 components. The first 20 BCR PCs were used to construct kNN graphs with k = 20 and to generate UMAP visualizations. If UMAP initialization failed because multiple cells had identical embeddings, UMAP was calculated for the unique embeddings and the resulting coordinates were assigned to cells sharing each embedding.

### BCR feature selection and encoding

BCR-derived features were matched to cells in the GEX Seurat object using cell barcodes. Evaluated features included isotype, SHM frequency, IGHV gene subgroup, clonal relationship, light chain locus identity (IGK or IGL), and CDRH3 features such as amino acid length. CDRH3 amino acid sequences were also represented using Atchley [72] and Kidera [73] physicochemical factors as implemented in immApex v1.4.4 [74] (Supplementary Figure 7).

BCR features were converted into numerical cell-level representations before integration with GEX. Categorical features, including isotype, were one-hot encoded with all observed categories retained. Continuous features, including SHM frequency and CDRH3 length, were represented directly. All features were standardized to a mean of zero and standard deviation of one.

### Integration of GEX and BCR data

BCR information was matched to the GEX Seurat object using cell barcodes. Paired light chain information was added to the corresponding heavy-chain records, and cells without both a heavy and light chain were excluded. The dataset was then restricted to cells annotated as B cells with paired BCR information, leaving 112,909 cells. For direct comparison of all five embedding methods, the data were further restricted to cells successfully processed by every model, leaving 111,713 cells. This filtering step also removed the small number of cells with BCRs with an IgE isotype.

### Weighted nearest-neighbor integration

GEX and BCR embeddings were integrated using the weighted nearest-neighbor (WNN) framework implemented in Seurat v4 [62]. FindMultiModalNeighbors was applied to the first 20 PCs from the GEX and BCR assays with k = 20. The default algorithm learned cell-specific modality weights based on within- and cross-modality predictions.

### Concatenation of GEX and BCR representations

We evaluated three approaches (*raw*, *reduced_gex*, and *reduced_both*) for concatenating GEX and BCR representations (either as language-model embeddings or standardized engineered features). In the *raw* approach, the original BCR embedding dimensions or features were combined with scaled expression values for the variable GEX features before PCA, neighbor detection, and UMAP calculation. In the *reduced_gex* approach, the BCR embedding dimensions or features were combined with the leading GEX PCs before dimensionality reduction and neighbor detection. In the *reduced_both* approach, PCA was run on the BCR representation, then BCR PCs were concatenated directly with GEX PCs before neighbor detection and UMAP calculation. Unless otherwise specified, 20 GEX PCs were used with 20 BCR PCs for embeddings or 5 BCR PCs for features (one per feature). Neighbor detection used k = 20.

For *reduced_gex* and *reduced_both* approaches, the modality-specific matrices were centered and rescaled before concatenation to contribute equal total variance. This prevented differences in dimensionality or scale from causing one modality to dominate the integrated space. The effects of unequal weighting were tested by varying the GEX weight from 0 to 1 (Supplementary Figure 11). For the main analyses, we used reduced_both for CE (40 total dimensions) and reduced_gex for CF (25 total dimensions); the other approaches for both spaces are compared in Supplementary Figure 12.

### Evaluation of cell-level spaces

All spaces were evaluated using local cell neighborhoods. Because Seurat can include a cell in its own kNN results, neighborhoods were initially calculated with k = 21. If the focal cell was present, it was removed; otherwise, the most distant neighbor was removed. This yielded 20 non-self neighbors for every cell in each space. Metrics were calculated both across the complete dataset and independently within each subject. For subject-level analyses, the Seurat object was divided by subject and neighbor graphs were recalculated within each subset, thereby preventing cross-subject neighbors from influencing the results.

### Categorical features

For each categorical feature, we counted the number of neighbors with the same value as the focal cell. Categories were consolidated when appropriate. For example, isotype was evaluated primarily as switched versus unswitched rather than as individual IgA, IgD, IgG, or IgM classes. A small number of cells (n = 1,486) were assigned both IGK and IGL light chain loci. These cells were treated as a distinct IGK, IGL category rather than as matching either locus individually. Clonal relationship agreement was evaluated only for cells that were part of expanded clonal families (i.e., clones containing at least two cells). To avoid penalizing small clones that could not occupy all 20 neighbor positions, the number of same-clone neighbors was divided by the smaller of k and the size of the clone minus one. Thus for example, if a cell was part of a clone of size 11 and 10 of its 20 nearest neighbors were clonally related, then we considered it to have 10/10 = 100% matching labels.

### Clustering and differential-expression analysis

Unsupervised clustering was performed separately for each space using Seurat FindClusters with the Leiden algorithm. To guide the selection of clustering resolution, mean silhouette scores were calculated across resolutions from 0.1 to 1.0 in increments of 0.1. Resolutions near the inflection point of each silhouette curve were selected.

A resolution of 0.4 was used for GEX, BCR, CE, and CF, and a resolution of 0.3 was used for GEX with IG genes retained. Silhouette scores were calculated using the first 20 PCs for the GEX and BCR spaces, all 40 CE PCs and all 25 CF PCs. Because WNN does not produce a directly corresponding joint PC space, its UMAP coordinates were used for this calculation. WNN produced many clusters even at low resolutions, and a resolution of 0.1 was therefore used. The selected resolutions produced 11 clusters for GEX, 10 for GEX with IG genes retained, 32 for BCR embeddings, 49 for WNN, 23 for CE, and 13 for CF. CellTypist was subsequently rerun with cluster-level majority voting enabled (majority_voting = TRUE).

Differentially expressed marker genes between clusters were evaluated using a pseudobulk strategy. Raw counts were aggregated for each cluster within each sample, and differential expression testing was performed with Seurat’s FindAllMarkers. Only genes with higher expression in the cluster (only.pos = TRUE), detected in at least 25% of cells in either group (min.pct = 0.25) and with log2 fold change ≥ 0.25 were tested. Genes with Bonferroni-adjusted P < 0.05 were considered differentially expressed.

### Statistics

All analyses were performed in R v4.5.2 [77]. Statistical significance was defined as nominal P values less than 0.05, except for differential expression where the Bonferroni-adjusted p value was used. Paired Wilcoxon signed-rank tests were used to compare subject-level neighborhood metrics between the GEX space and each alternative space.

### Code and data availability

All analysis code is available at https://github.com/Kleinstein-Lab/athanor and https://github.com/Kleinstein-Lab/gex-bcr-integration. Public data were taken from GSE270142 (Diabetes Nicholas) and GSE193869 (Lupus Siu).

## Author contributions

SHK conceptualized the study, EA performed data acquisition and analysis with all authors participating in interpretation. EA wrote the manuscript with all authors editing and approving the final content for submission.

## Acknowledgments

This work was supported by the National Institute of Allergy and Infectious Diseases of the NIH through the R01AI104739 and U19AI089992 and by the National Library of Medicine through the T15LM007056. It is the result of NIH funding, in whole or in part, and is subject to the NIH Public Access Policy. Through acceptance of this federal funding, the NIH has been given a right to make the work publicly available in PubMed Central. The content is solely the responsibility of the authors and does not necessarily represent the official views of the NIH.

We thank Meng Wang for advice on BCR embeddings and Yuval Kluger, Edward Lee, Cole Jensen, and Dylan Duchen for advice on integration and insights into B cell biology. We also thank Linda Bockenstedt for reviewing the manuscript draft.

## Footnotes

### Confficts of interest

SHK receives consulting fees from Peraton.

