## Supplementary Methods for "Cell-level integration of gene expression and B cell receptor data captures complementary B cell information"

Unless otherwise specified, processing steps are the same as in Methods.

*Data collection and sample processing*

The Lupus Siu dataset had 15 samples from nine subjects; three subjects (A, B, and C) each had samples taken from three tissues (appendix, lymph node, and spleen), three subjects had systemic lupus erythematosus (SLE 1, 2 and 3) and three subjects were healthy control donors (HCD 1, 2, and 3). The SLE and HCD donor samples were taken from PBMCs. All samples were sorted using $CD19^+$ and B cells were labelled by the original authors as activated but not classical B cells, activated naïve B cells, double negative ($CD27^-IgD^-$) B cells, germinal center B cells, IgM only B cells, memory B cells, marginal zone B cells, naïve B cells, plasmablasts, and transitional B cells.

*GEX data processing*

There were 103,531 cells initially available, 100,980 cells after quality control, and 91,846 cells after doublet removal (prior to subsetting down to just B cells).

Batch effects were assessed by inspecting UMAPs colored by subject. Since subject-driven structure was evident, Harmony integration (group.by = subject) was applied to the PCA reduction, and neighbors and UMAPs were recomputed on the integrated (harmony) reduction using the same number of dimensions and neighbors. The integrated neighbors were used for all downstream calculations, including the integration with BCR embeddings or features and WNN.

*ADT data processing*

In addition to CD21, CD27, IgD and IgM, we also used CD38 as an ADT marker since it was available in this dataset.

*BCR data processing*

Since the AIRR rearrangement files were unavailable for this dataset, they were generated from Cell Ranger V(D)J outputs (filtered_contig.fasta and filtered_contig_annotations.csv) using the Immcantation v4.5.0 changeo-10x pipeline.

*Integration of GEX and BCR data*

After removing cells without paired BCRs and non-B cells, there were 50,908 cells left. The final cell count after subsetting down to cells with BCRs that ran through all five embedding methods was 49,851 cells.

*Clustering*

Clustering resolutions were chosen the same way as for the Diabetes Nicholas data (the inflection point of the silhouette score plot) but the resolutions differed. We chose 0.3 for GEX (8 clusters) and 0.7 for CF (16 clusters).
