## Supplementary Figures for "Cell-level integration of gene expression and B cell receptor data captures complementary B cell information"

*Supplementary Figure 1 (Clustering resolution range)*

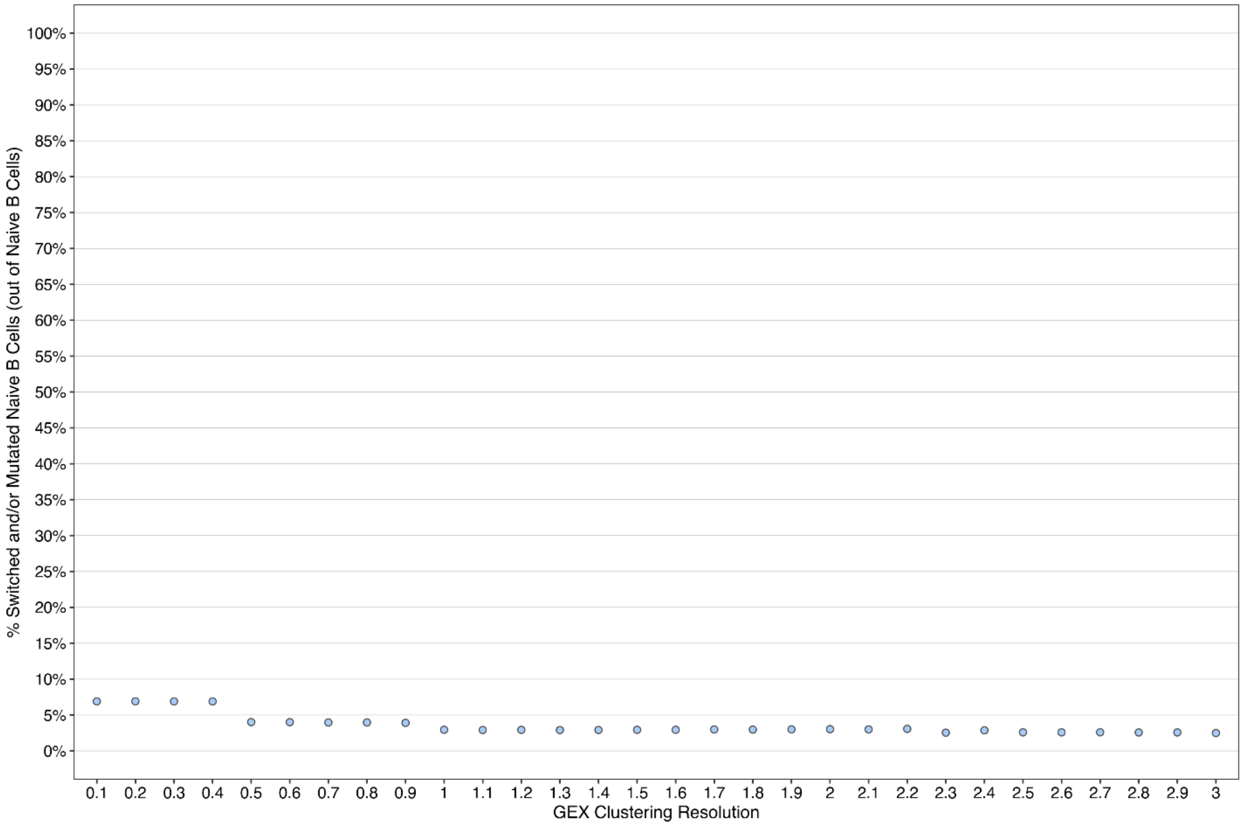


***Supplementary Figure 1. The frequency of cells annotated as naïve B cells that express BCRs that are class-switched and/or mutated is not strongly impacted by clustering resolution.*** Percentage of the total naïve B cells that were switched and/or mutated after clustering the GEX space at resolutions from 0.1 to 3. Cell type annotation was done using CellTypist with majority voting and the Adult Human Blood reference.

*Supplementary Figure 2 (Cell level GEX)*

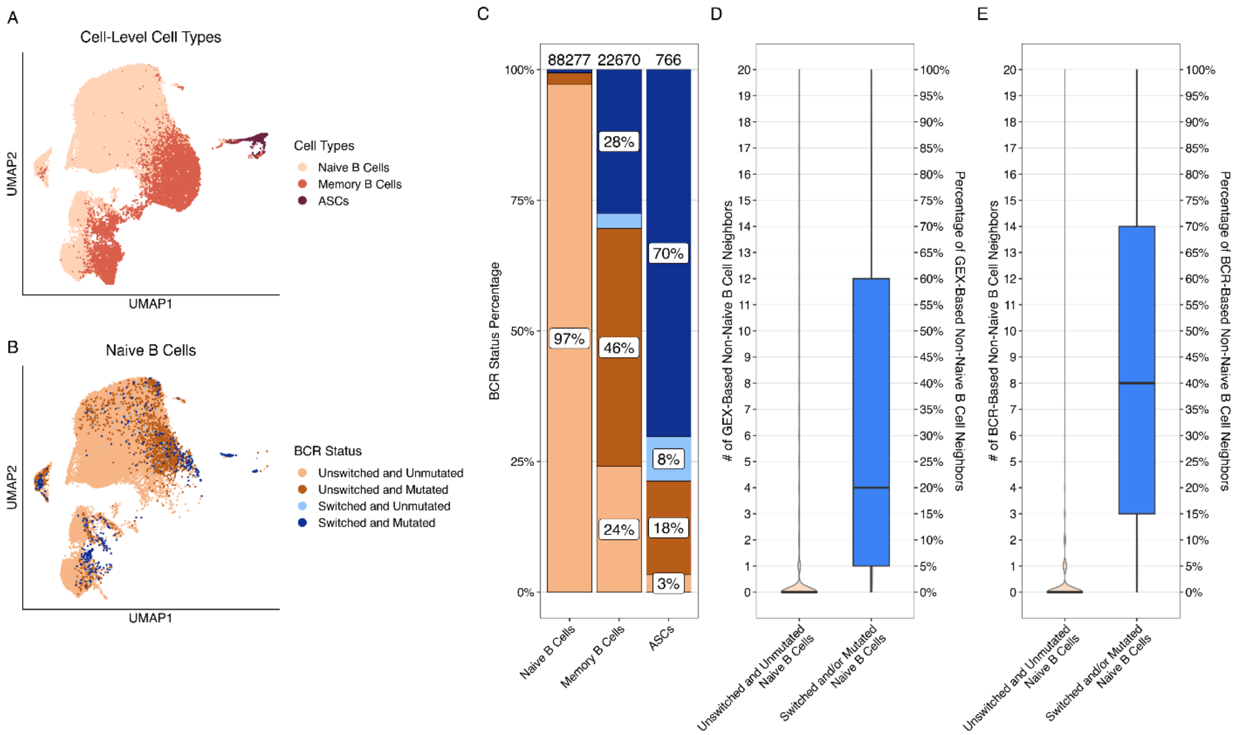


***Supplementary Figure 2. BCR features are inconsistent with individual cell-based B cell annotations.*** **A)** UMAP of the GEX space, colored by cell types, as annotated by CellTypist on a cell-level with the Adult Human Blood reference. **B)** The same UMAP restricted to naïve B cells and colored by BCR status. **C)** Stacked bar plot showing the composition of each cell type by BCR status (colored the same as B). Total cell counts for each cell type are shown above each bar. **D)** Box plot of the number (left axis) and percentage (right axis) of each cell's k = 20 nearest GEX-defined non-naïve B cell neighbors, compared by the unswitched and unmutated naïve B cells vs. the switched and/or mutated naïve B cells. Violin plots behind each box plot show the full distribution of values. Boxes span the interquartile range (IQR) with the median line; whiskers extend to 1.5x IQR. **E)** The same comparison as (D) but quantified over BCR-defined non-naïve B cell neighbors.

*Supplementary Figure 3 (Effects of different k's)*

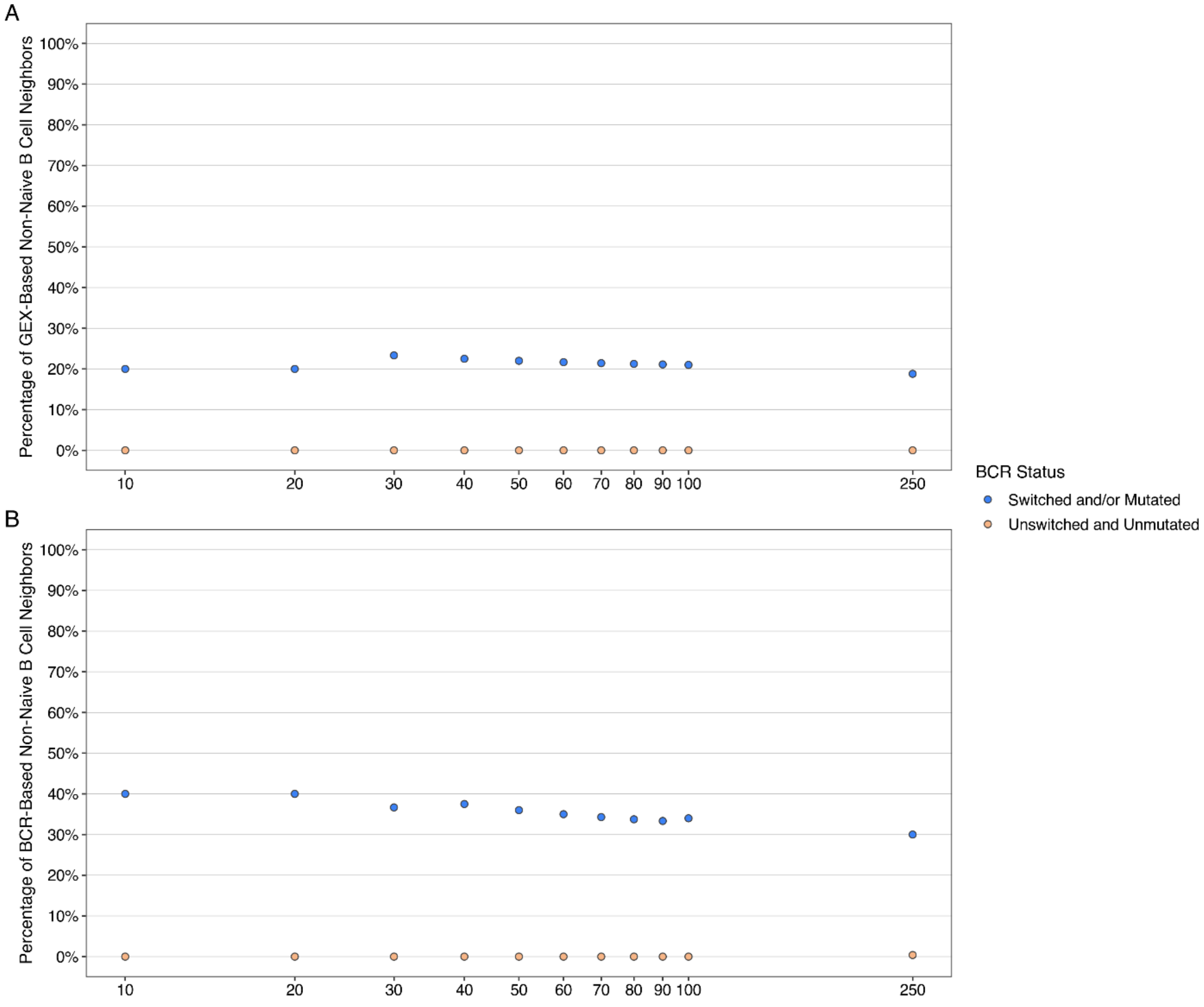


***Supplementary Figure 3. The consistency of the local neighborhood of correctly annotated and misannotated naïve B cells is robust to the choice of k.*** **A)** Median percentage of each naïve B cell's k-nearest neighbors that were defined as non-naïve B cells using GEX. k ranged from 10 to 100 by 10s, with 250 included as an example of an even larger k. Naïve B cells were split by BCR status into unswitched and unmutated naïve B cells (orange) and switched and/or mutated naïve B cells (blue). **B)** The same as (A) but for BCR-defined non-naïve B cells. Naïve B cells were labelled at the cell level.

*Supplementary Figure 4 (BCR embedding choice)*

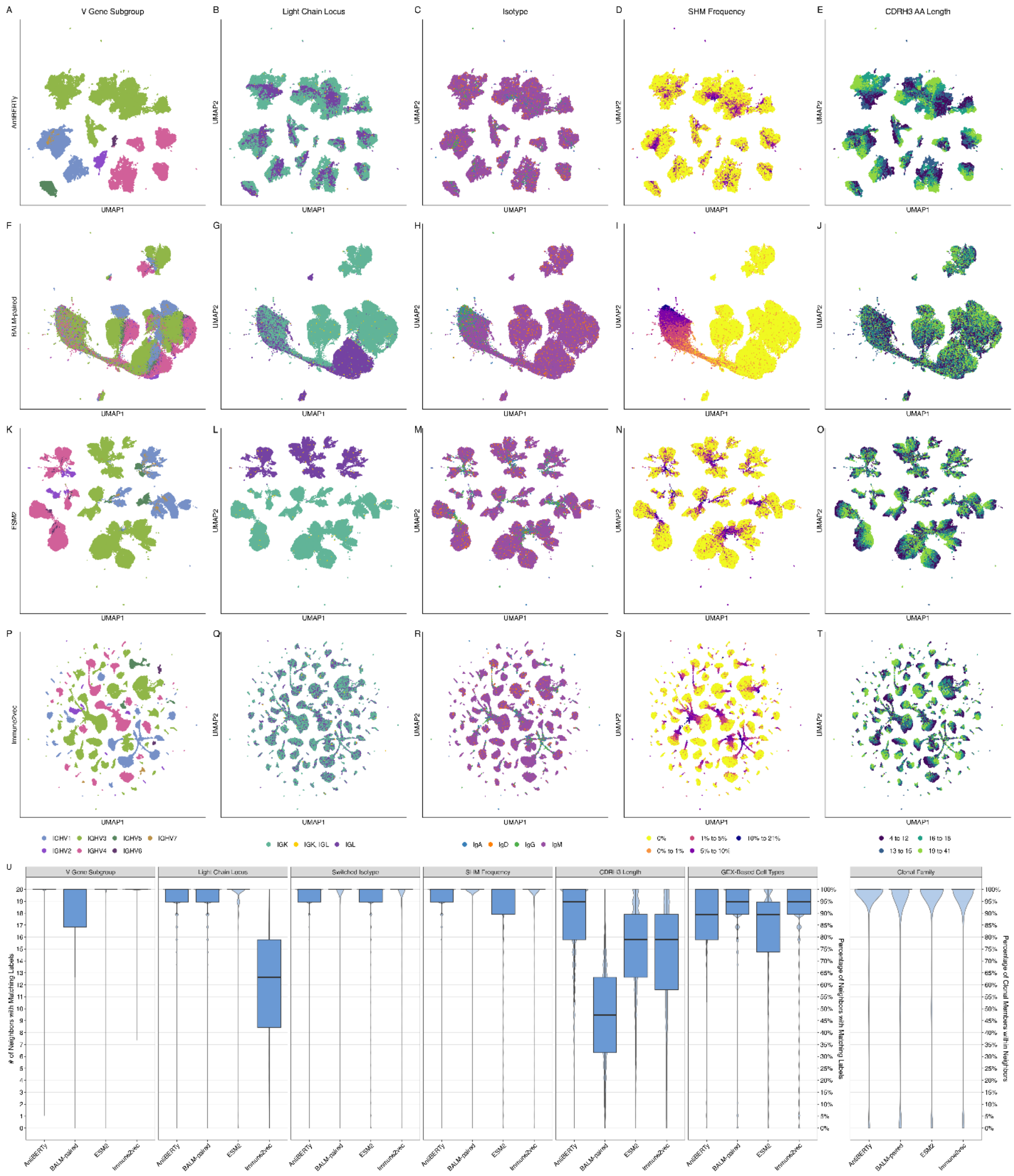


***Supplementary Figure 4. BCR features are preserved across embedding methods.*** **A-T)** UMAPs of BCR spaces generated using four different language models: AntiBERTy (A-E), BALM-paired (F-J), ESM2 (K-O), and Immune2vec (P-T). Columns are colored by BCR features, from left to right: V gene subgroup, light chain locus, isotype, binned SHM frequency, and binned CDRH3 amino acid length. **U)** Box plots showing the number (left axis) and percentage (right axis) of each cell's k = 20 nearest neighbors that share the same BCR feature (V gene subgroup, light chain locus, isotype, SHM frequency, CDRH3 length within 2 amino acids) or GEX-based cell type annotation as indicated on each panel, for each embedding method. Clonal relationship matches (right) were calculated for expanded clones only, with percentages calculated out of the minimum relative to the smaller of the clone size and k. Violin plots behind each box plot show the full distribution of values. Boxes span the IQR with the median and whiskers extend to 1.5x IQR.

*Supplementary Figure 5 (Figure 2 extended)*

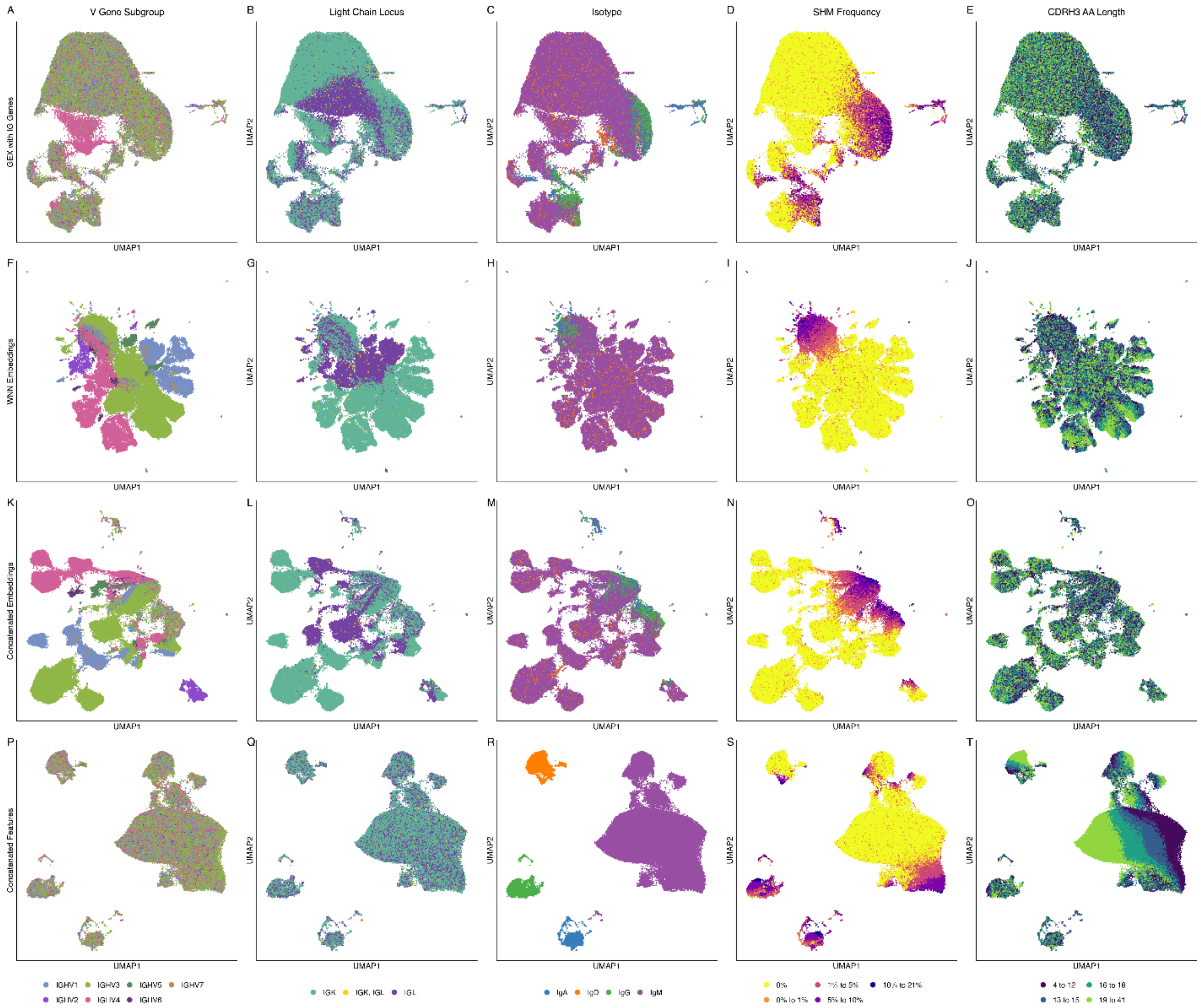


***Supplementary Figure 5. BCR features in GEX+IG and integrated spaces.*** UMAPs of GEX with IG genes retained **(A-E)**, WNN embeddings **(F-J)**, concatenated embeddings **(K-O)**, and concatenated features **(P-T)**. Columns are colored by BCR features, from left to right: V gene subgroup, light chain locus, isotype, binned SHM frequency, and binned CDRH3 amino acid length.

*Supplementary Figure 6 (CDRH3 length ranges for neighbor matching)*

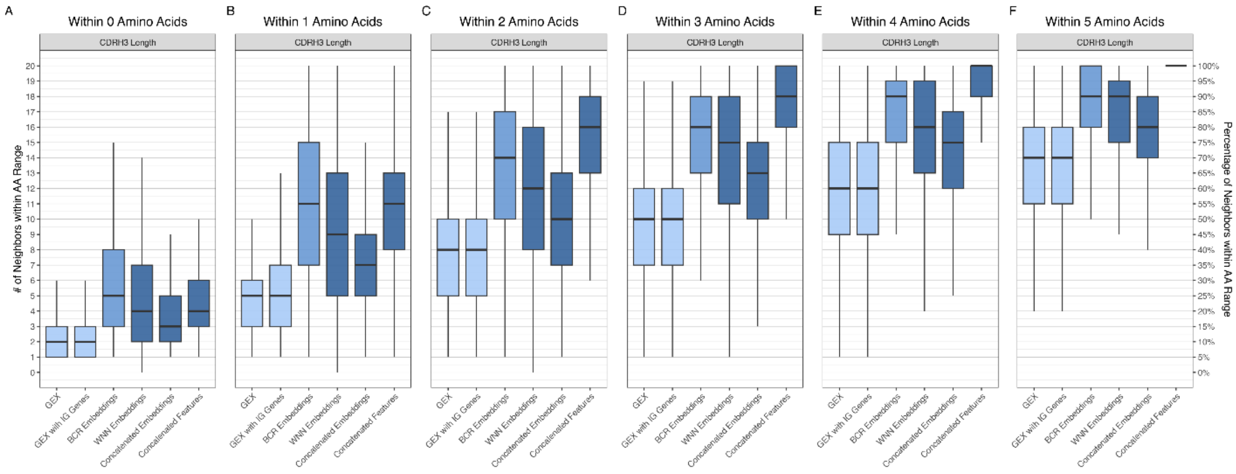


***Supplementary Figure 6. Local neighborhood consistency of CDRH3 length at increasing tolerances.* A-F)** Box plots showing the number (left axis) and percentage (right axis) of each cell's k = 20 nearest neighbors whose CDRH3 amino acid length is within 0 (A) to 5 (F) amino acids of the cell's own CDRH3 length, for each of the six spaces. Fill color indicates the data used to build each space: GEX (light blue), BCR (medium blue), or GEX and BCR (dark blue). Boxes span the IQR with the median and whiskers extend to 1.5x IQR.

*Supplementary Figure 7 (Physicochemical correlations)*

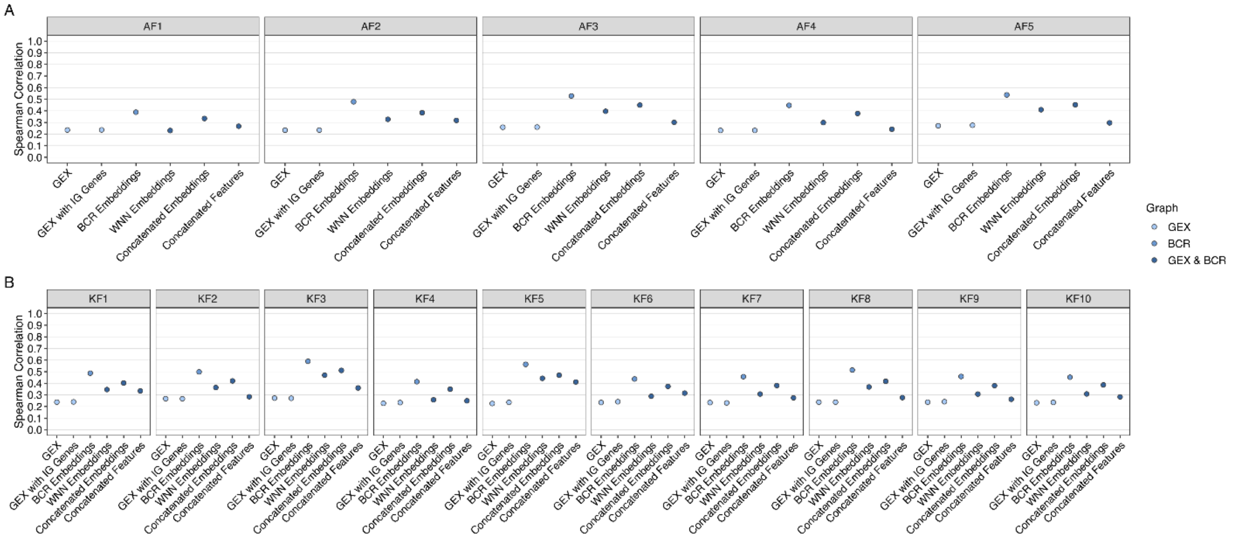


***Supplementary Figure 7. Spearman correlations of CDRH3 physicochemical properties across spaces.*** Spearman correlation between each cell's **A)** Atchley factor (AF1 to AF5) or **B)** Kidera factor (KF1 to KF10) and the mean value among its k = 20 nearest neighbors, calculated across all cells for each of the six spaces.

*Supplementary Figure 8 (ADT correlation alternative)*

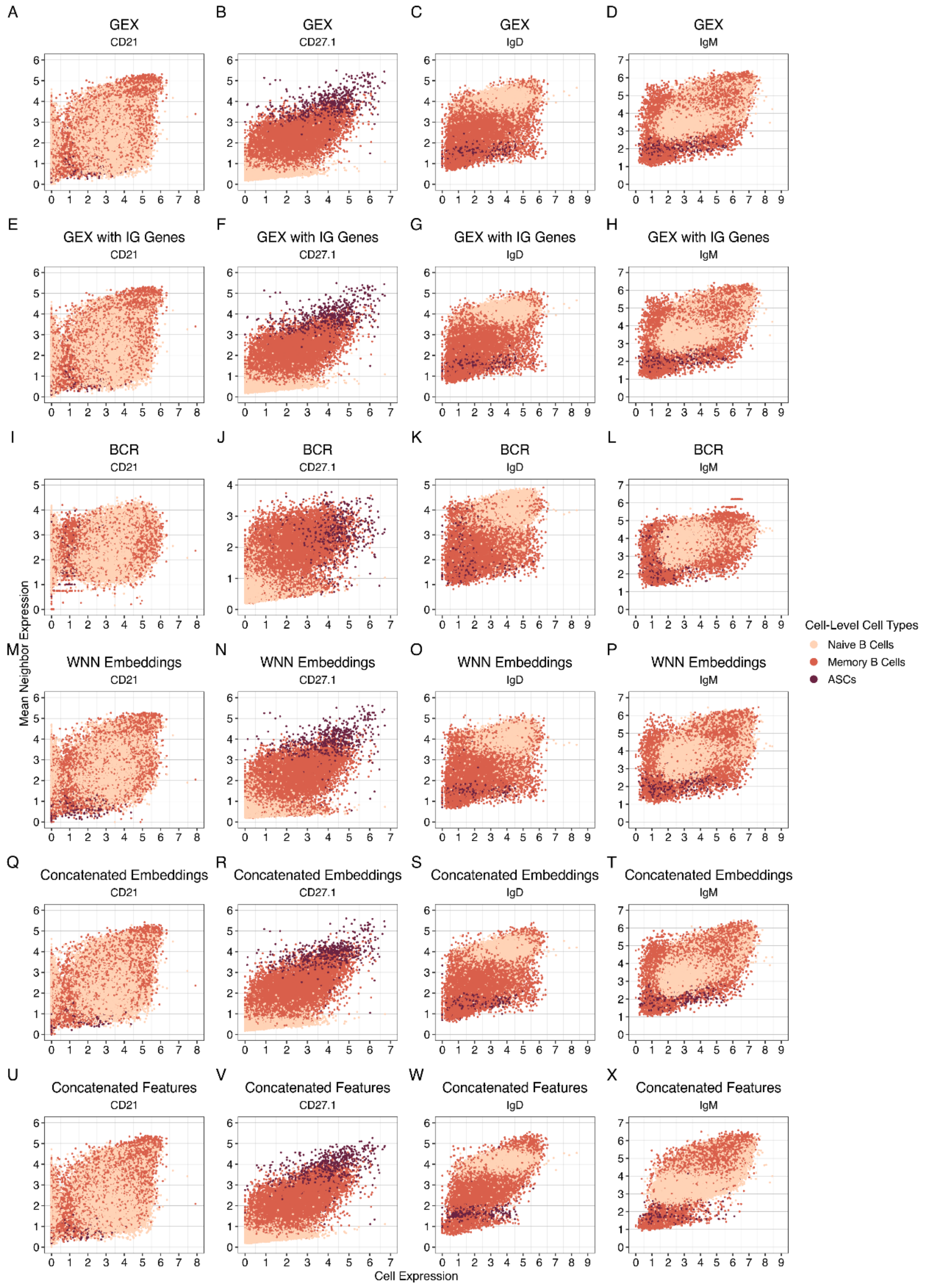


***Supplementary Figure 8. Relationships between each cell's surface protein expression and its neighbors.* A-X)** Scatter plots of each cell's expression of four B cell

ADT markers on the x-axis (CD21, CD27.1, IgD, IgM) against the mean expression among its k = 20 nearest neighbors on the y-axis, for each of the six spaces (rows). Cells are colored by cell-level cell type annotation.

*Supplementary Figure 9 (Evaluation metrics)*

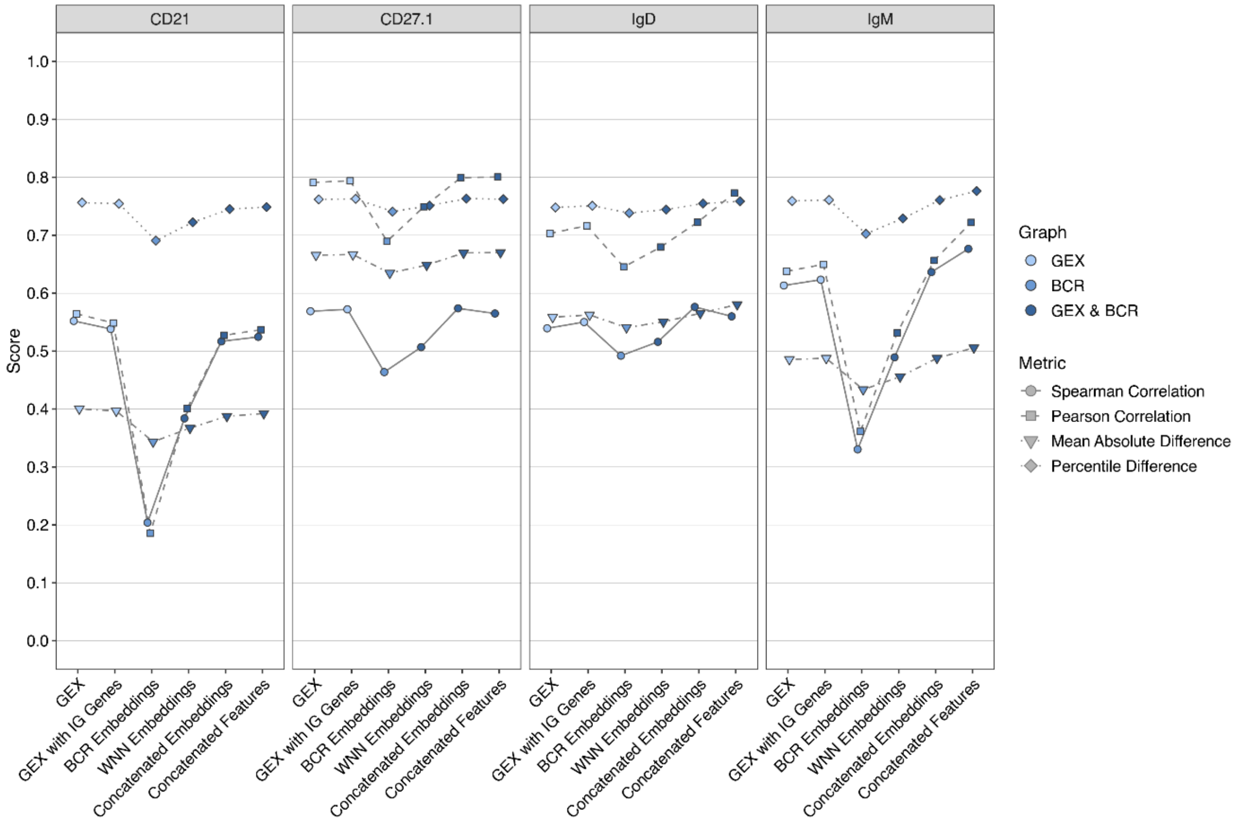


***Supplementary Figure 9. Comparison of evaluation metrics for local neighborhood consistency of cell surface protein expression.*** For each of the six spaces (x-axis), we evaluated how well each cell's expression of four B cell ADT markers (CD21, CD27.1, IgD, IgM) relates to its k = 20 nearest neighbors. Four metrics were computed: Spearman correlation, Pearson correlation, mean absolute difference (MAD), and percentile difference (PD). Spearman (circular points, solid lines) and Pearson (square points, dashed lines) correlations compare each cell's expression with the mean expression among its neighbors, calculated across all cells. MAD (triangular points, dot-dashed lines) is the absolute difference between each cell's expression and its neighbors' mean expression, averaged across all cells. PD (diamond points, dotted lines) is computed the same way, but with each cell's expression converted to a percentile rank to make it less sensitive to each marker's scale. To place all metrics onto a common 0-1 scale with higher values indicating better neighborhood consistency, MAD was transformed as 1 / (1 + MAD) and PD as 1 - PD. Fill color indicates the data used to build each space: GEX (light blue), BCR (medium blue), or GEX and BCR (dark blue).

*Supplementary Figure 10 (AMULETY constant regions)*

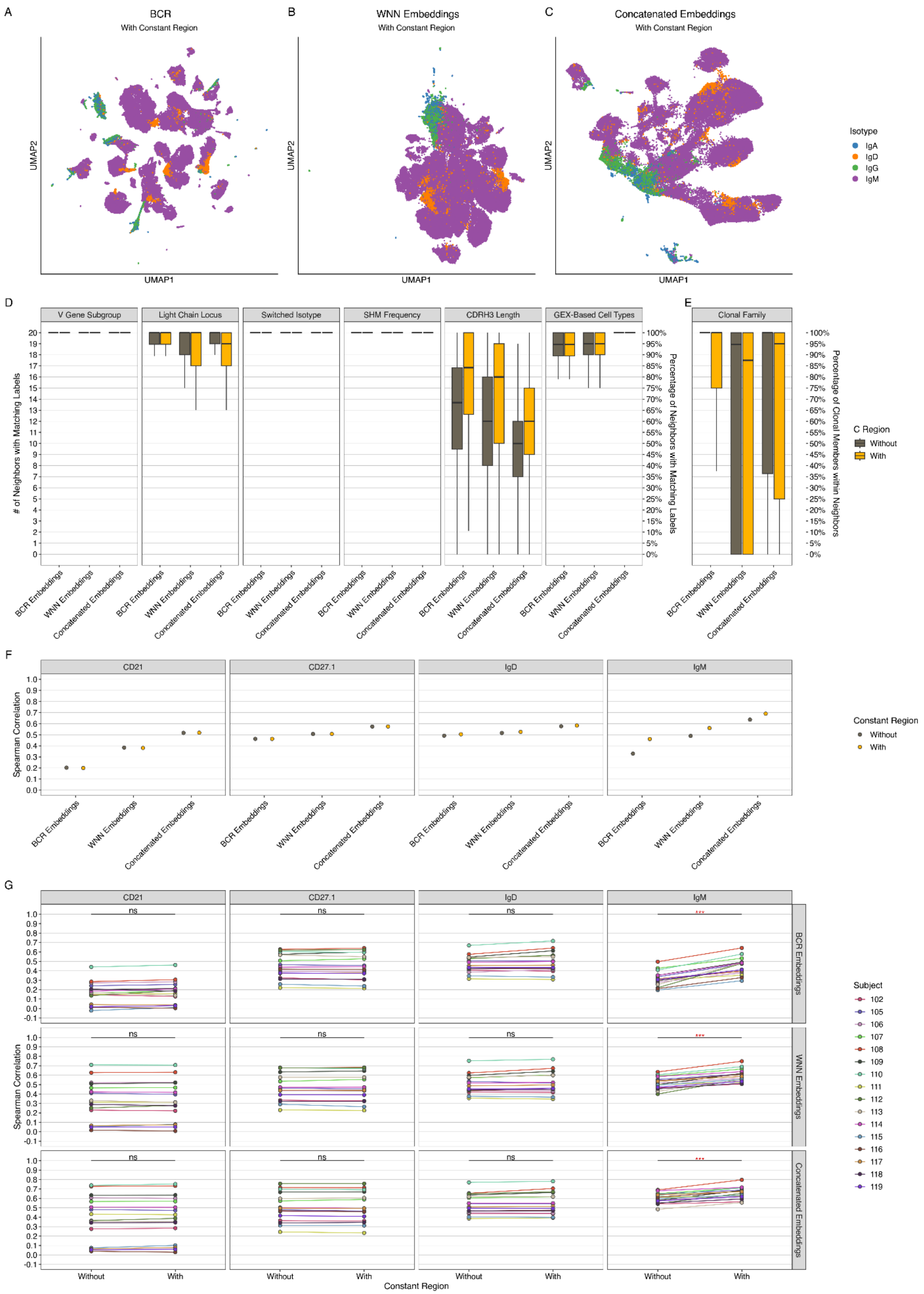

***Supplementary Figure 10. Effect of including the constant region in BCR embeddings.***
**A-C)** UMAPs of (A) BCR, (B) WNN embeddings, and (C) concatenated embeddings spaces generated with the constant region included, colored by isotype. **D-E)** Box plots showing the number of each cell's k = 20 nearest neighbors that share the same BCR feature, GEX-based cell type annotation, or clonal family, for spaces generated without (gray) or with (yellow) the constant region. Boxes span the IQR with the median and whiskers extend to 1.5x IQR. **F)** Spearman correlation between each cell's expression of four B cell ADT markers (CD21, CD27.1, IgD, IgM) and the mean expression among its k = 20 nearest neighbors, calculated across all cells. **G)** Per-subject Spearman correlations for the four ADT markers in (F). Lines connect values from the same subject. Within each space, spaces with and without the constant region were compared using a paired Wilcoxon signed-rank test across subjects (***P < 0.001; **P < 0.01; *P < 0.05; ns, not significant). Red asterisks indicate significant increases with the constant region.

*Supplementary Figure 11 (Effects of different modality weights)*

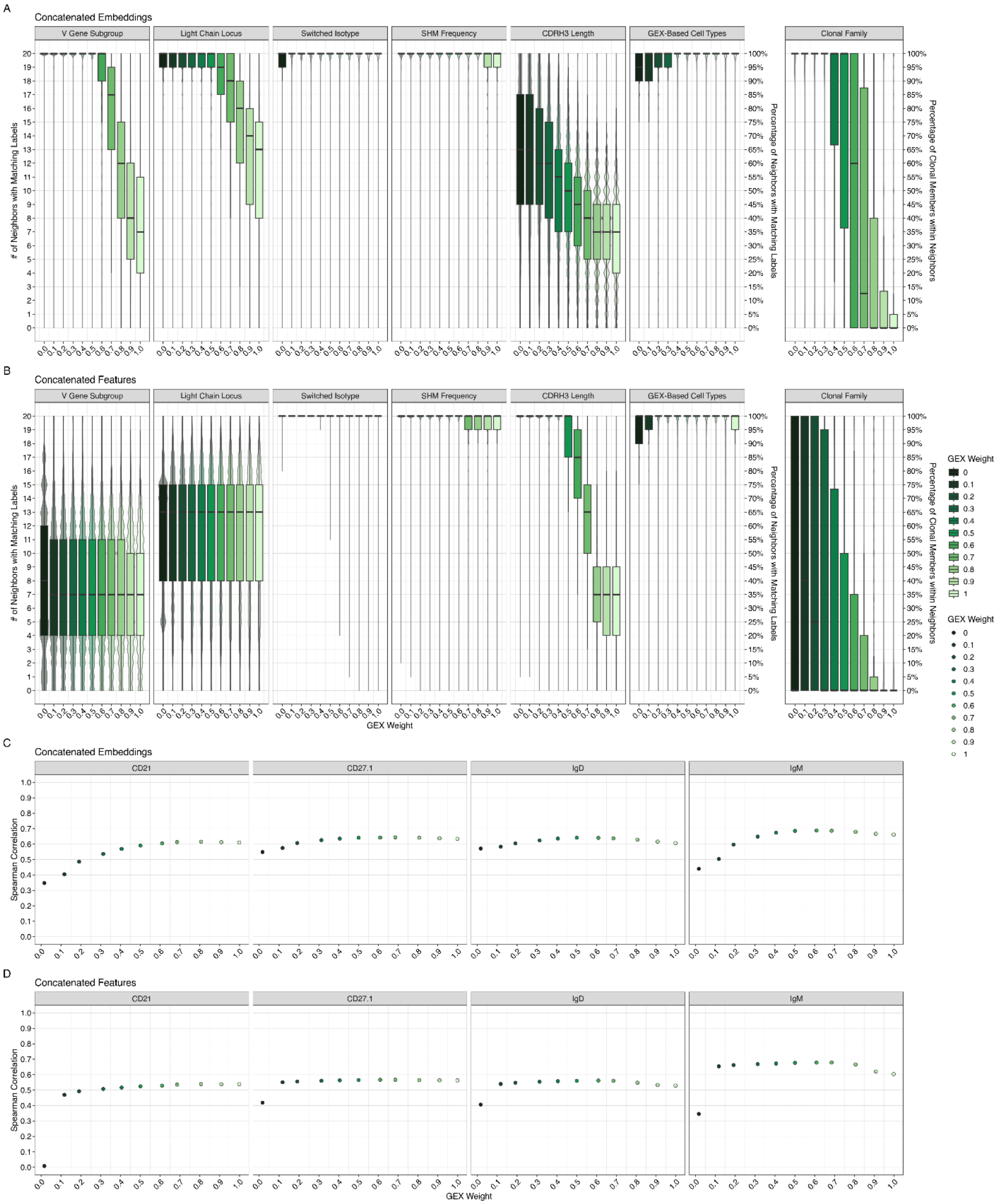


***Supplementary Figure 11. Effects of different modality weights on concatenated spaces. A-B)*** Box plots showing the number (left axis) and percentage (right axis) of each cell's k = 20 nearest neighbors that share the same BCR feature (V gene subgroup, light chain locus, isotype, SHM frequency, CDRH3 length within 2 amino acids) or GEX-based cell type annotation, for (A) concatenated embeddings and (B) concatenated features. The GEX weights range from 0 (BCR only) to 1 (GEX only). Violin plots behind each box plot show the full distribution of values. Boxes span the IQR with the median and whiskers extend to 1.5x IQR. **C-D)** Spearman correlation between each cell's expression of four B cell ADT markers (CD21, CD27.1, IgD, IgM) and the mean expression among its k = 20 nearest neighbors, across the same GEX weights for (C) concatenated embeddings and (D) concatenated features.

*Supplementary Figure 12 (Concatenation approach comparison)*

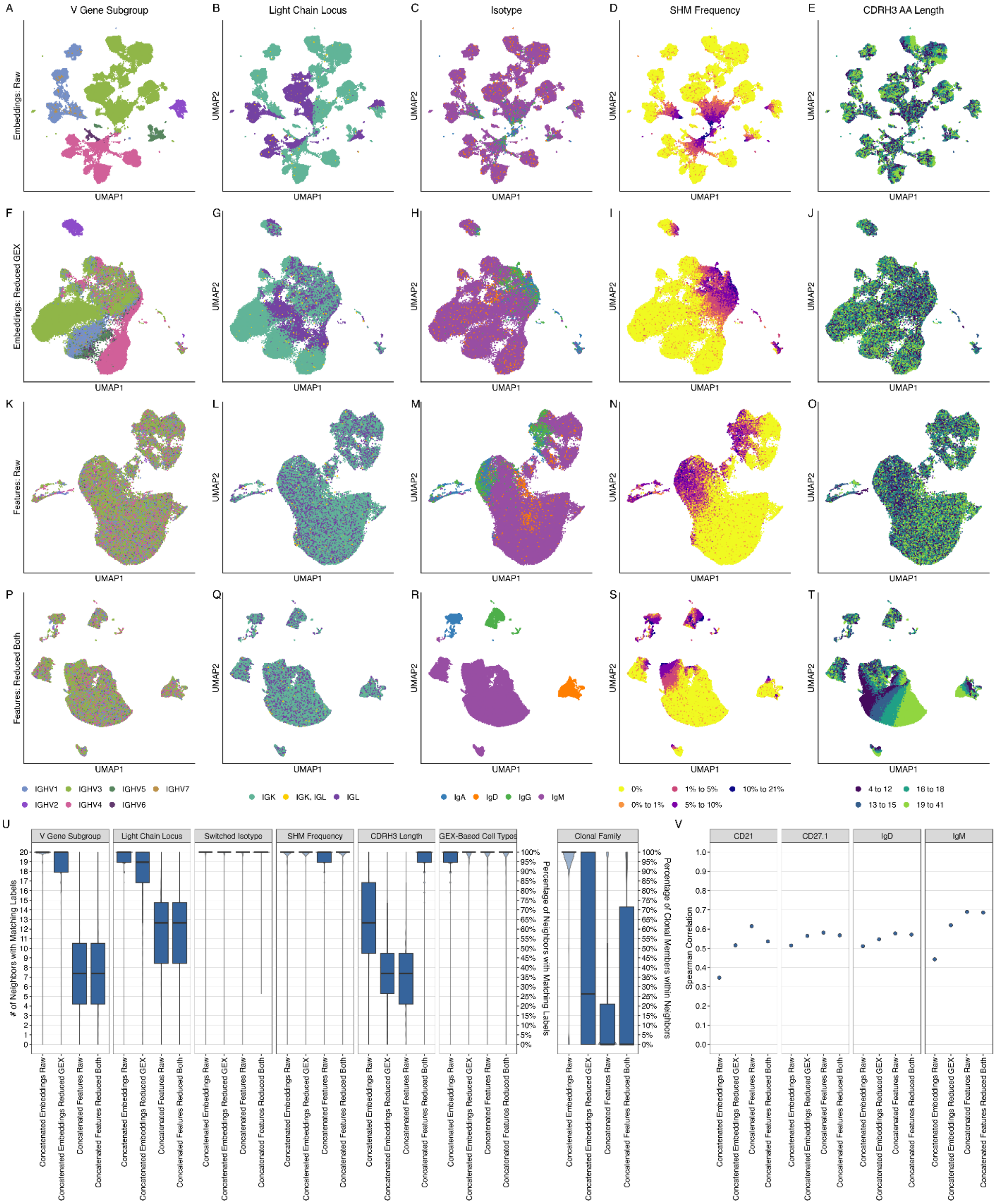


***Supplementary Figure 12. Comparison of concatenation approaches.*** **A-T)** UMAPs of concatenated spaces built from "raw" embeddings (A-E), "reduced_gex" embeddings (F-J), "raw" features (K-O), and "reduced_both" features (P-T). Each column is colored by a BCR feature, from left to right: V gene subgroup, light chain locus, isotype, binned SHM frequency, and binned CDRH3 amino acid length. **U)** Box plots showing the number (left axis) and percentage (right axis) of each cell's k = 20 nearest neighbors that share the same BCR feature or GEX-based cell type annotation, for each approach. Violin plots behind each box plot show the full distribution of values. Boxes span the IQR with the median and whiskers extend to 1.5x IQR. **V)** Spearman correlation between each cell's expression of four B cell ADT markers (CD21, CD27.1, IgD, IgM) and the mean expression among its k = 20 nearest neighbors, for each approach.

*Supplementary Figure 13 (Effects of different nPCs)*

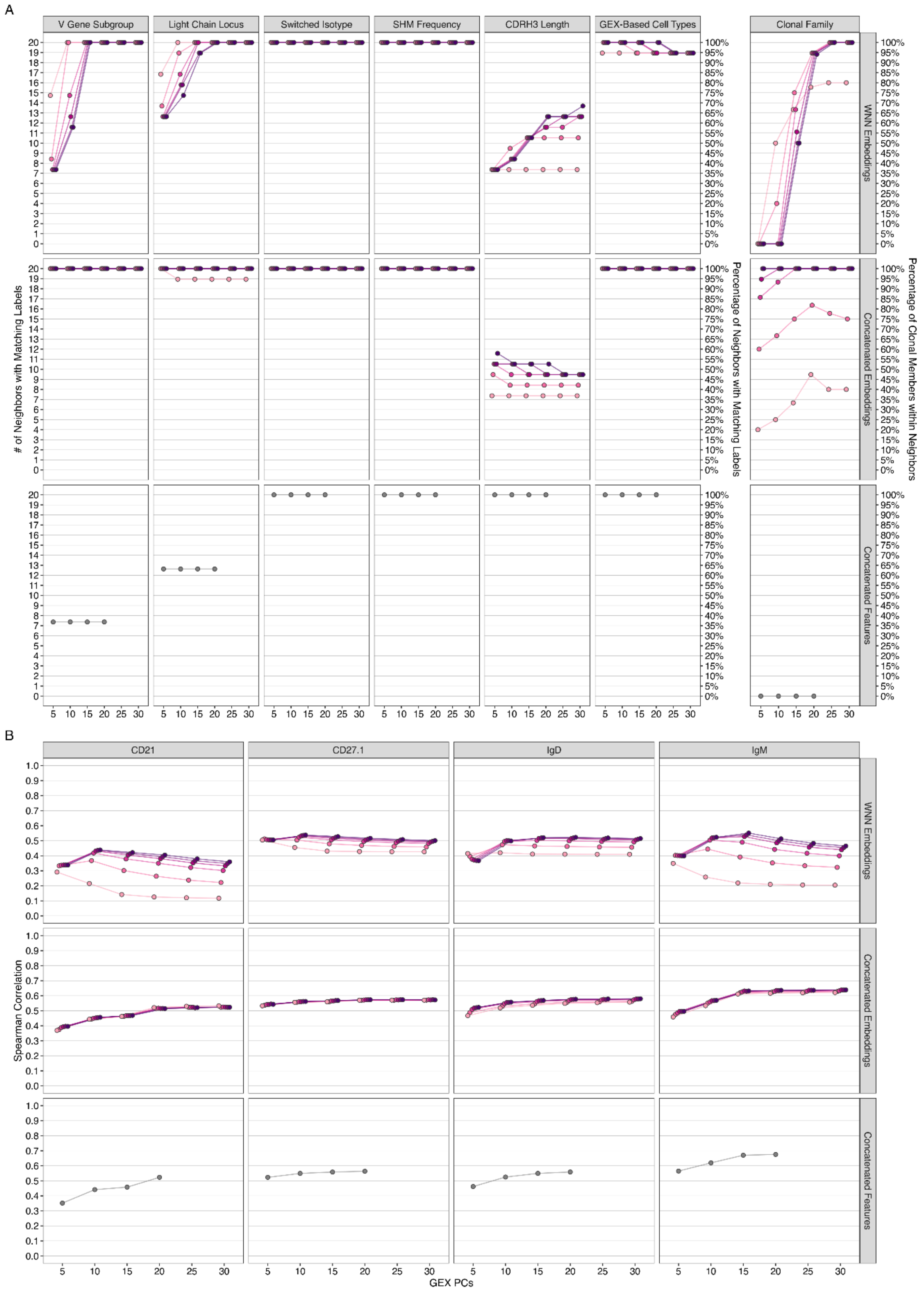

***Supplementary Figure 13. Effect of the number of principal components on integrated spaces.*** **A)** Median number (left axis) and percentage (right axis) of each cell's k = 20 nearest neighbors that share the same BCR feature, GEX-based cell type annotation, or clonal family, for WNN embeddings, concatenated embeddings, and concatenated features spaces built with varying numbers of GEX PCs (x-axis) and BCR PCs (color). Concatenated features do not use BCR PCs (gray). **B)** Spearman correlation between each cell's expression of four B cell ADT markers (CD21, CD27.1, IgD, IgM) and the mean expression among its k = 20 nearest neighbors, for the same spaces and PC combinations.

*Supplementary Figure 14 (Non-naïve neighbors)*

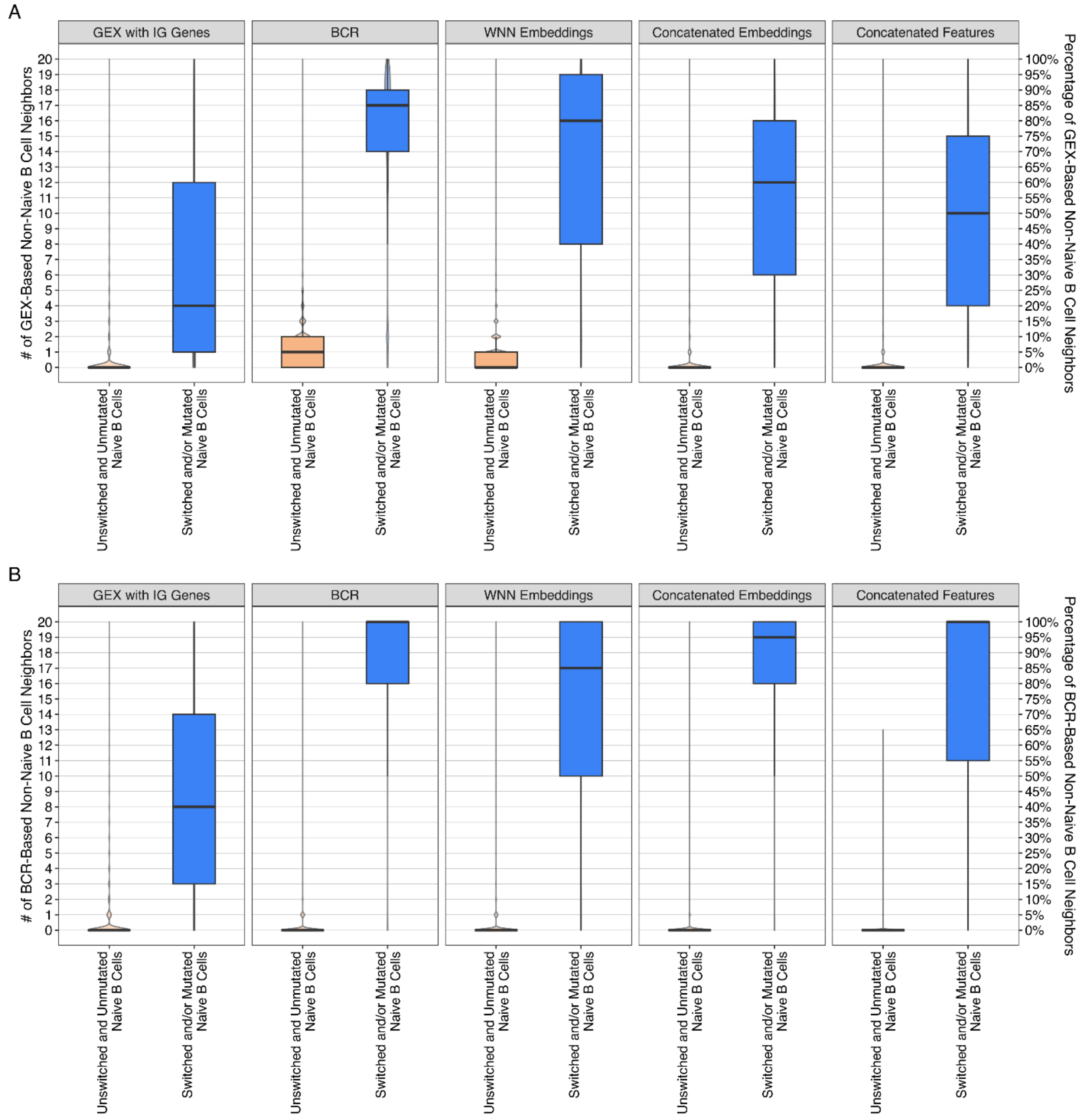


***Supplementary Figure 14. Local neighborhood consistency of correctly annotated and misannotated naïve B cells across spaces.*** **A)** Number (left axis) and percentage (right axis) of each naïve B cell's k = 20 nearest neighbors that were annotated as non-naïve B cells, for GEX with IG genes, BCR, WNN embeddings, concatenated embeddings, and concatenated features spaces. Naïve B cells are grouped by BCR status as unswitched and unmutated (left) or switched and/or mutated (right) and labeled at the cell-level using CellTypist with the Adult Human Blood reference. **B)** Same as in (A), but for neighbors with non-naïve BCR characteristics (switched and/or mutated). Violin plots behind each box plot show the full distribution of values. Boxes span the IQR with the median and whiskers extend to 1.5x IQR.

*Supplementary Figure 15 (Other cell type annotation references)*

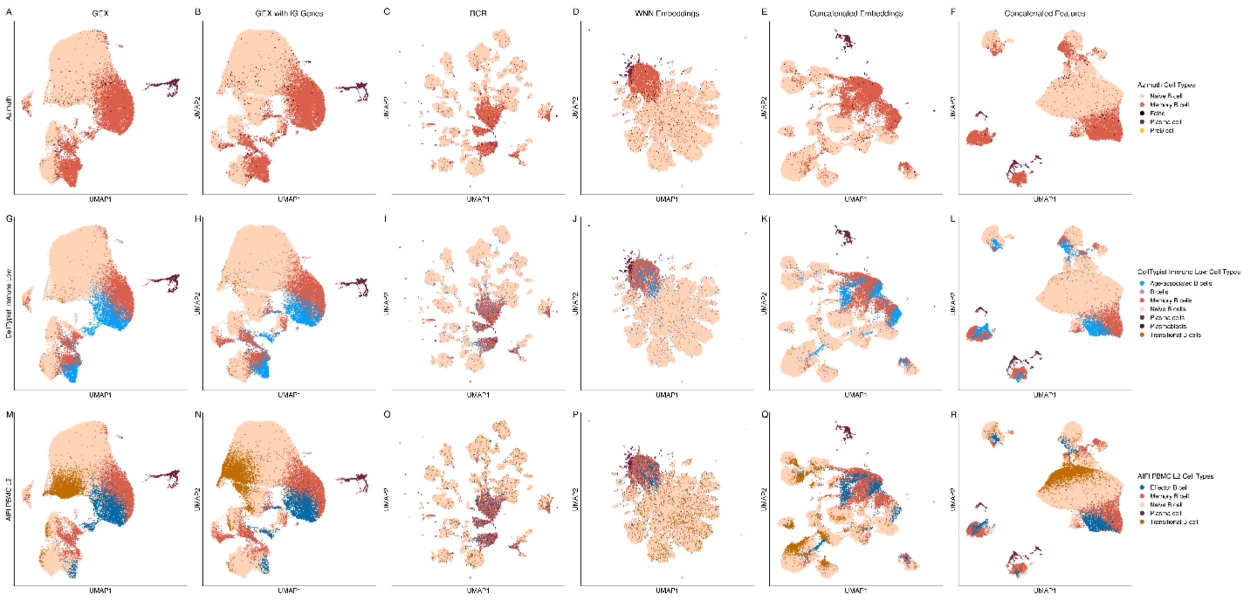


***Supplementary Figure 15. Cell-level annotations with alternative cell type references.*** UMAPs of each of the six spaces (columns), colored by cell-level cell type annotations from three alternative references: Azimuth **(A-F)**, CellTypist Immune All Low **(G-L)**, and the Allen Institute for Immunology's PBMC B cell reference at level 2 **(M-R)**.

*Supplementary Figure 16 (Lupus Siu)*

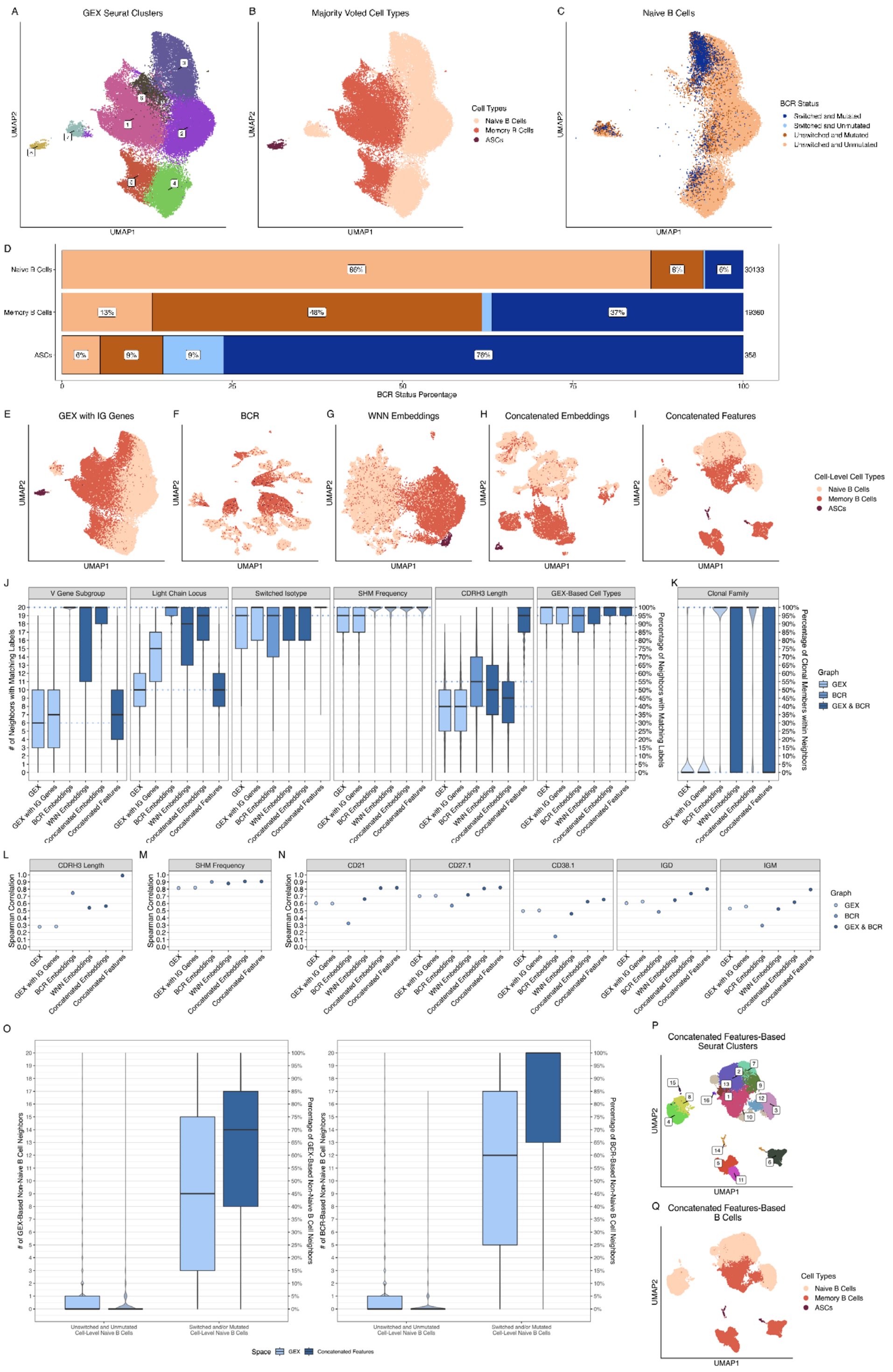


***Supplementary Figure 16. Validation in the Lupus Siu dataset.* A)** UMAP of the GEX space, colored by the 8 clusters identified by Leiden clustering (resolution = 0.3). **B)** The same UMAP colored by cell types, as annotated by CellTypist with cluster majority voting.

**C)** The same UMAP restricted to naïve B cells and colored by BCR status. **D)** Stacked bar plot showing the composition of each cell type by BCR status (colored as in C). Total cell counts for each cell type are shown to the right of each bar. **E-I)** UMAPs of (E) GEX with IG genes, (F) BCR embeddings, (G) WNN embeddings, (H) concatenated embeddings, and (I) concatenated features spaces, colored by cell-level cell type annotations. **J)** Box plots showing the number (left axis) and percentage (right axis) of each cell's k = 20 nearest neighbors that share the same BCR feature or GEX-based cell type annotation, for each space. Dotted lines indicate the medians for GEX (light blue) and BCR embeddings (medium blue). **K)** Box plot calculated as in (J), but for clonal relationships among cells (for expanded clones only), with percentages calculated relative to the smaller of the clone size and k. Violin plots behind each box plot show the full distribution of values. Boxes span the IQR with the median and whiskers extend to 1.5x IQR. **L-M)** Spearman correlation between each cell's (L) CDRH3 amino acid length or (M) SHM frequency and the mean value among its k = 20 nearest neighbors, for each space. **N)** Spearman correlation between each cell's expression of five B cell ADT markers (CD21, CD27.1, CD38.1, IgD, IgM) and the mean expression among its k = 20 nearest neighbors. **O)** Number (left axis) and percentage (right axis) of each cell-level naïve B cell's k = 20 nearest neighbors in the GEX (light blue) and CF (dark blue) spaces that were annotated as non-naïve B cells (left) or had non-naïve BCR characteristics (right). Naïve B cells are grouped by BCR status. **P)** UMAP of the CF space, colored by the 16 clusters identified by Leiden clustering (resolution = 0.7). **Q)** The same UMAP colored by cell type, assigned by CellTypist with cluster majority voting.

*Supplementary Figure 17 (Workflow overview)*

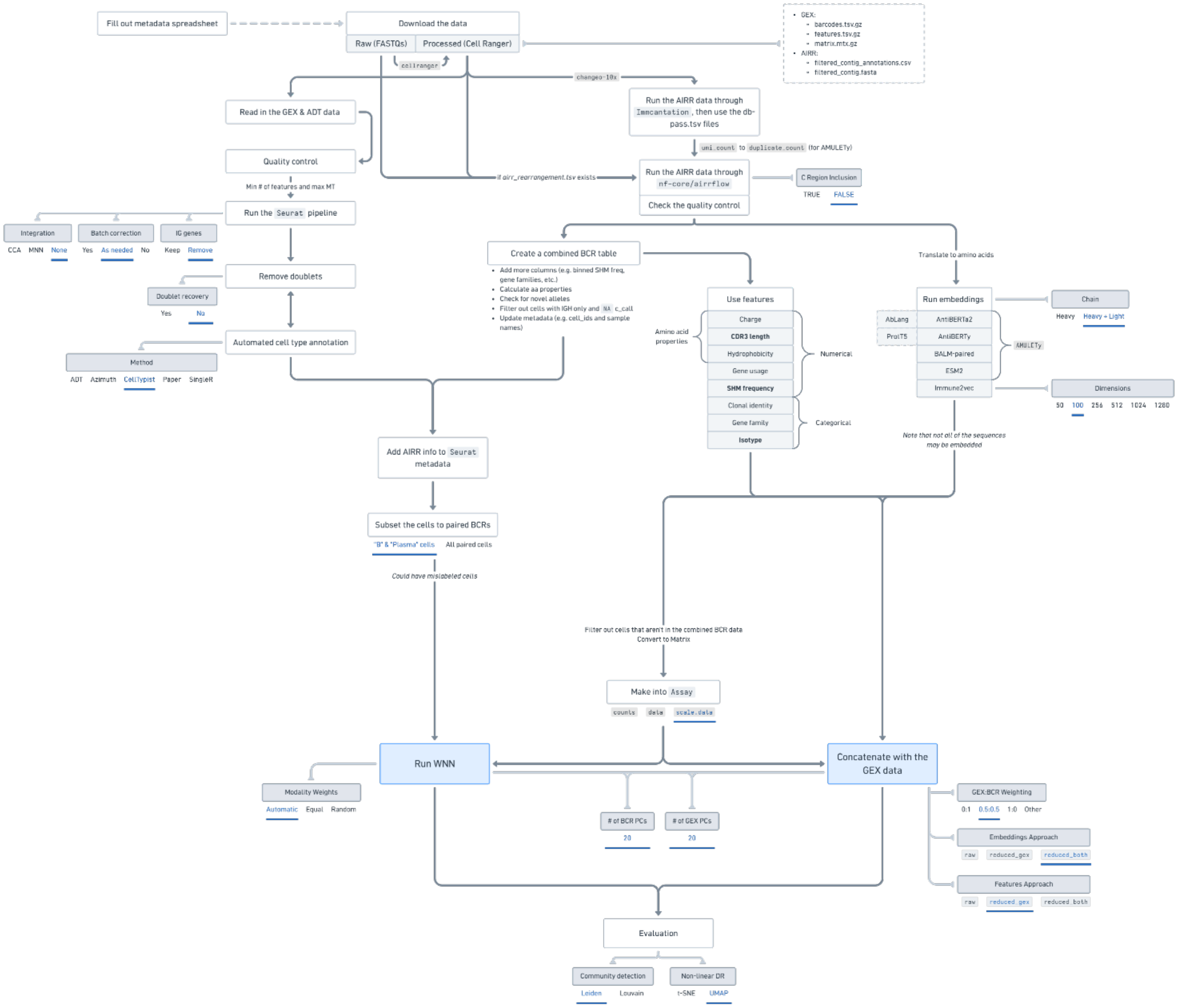


***Supplementary Figure 17. Overview of the analysis workflow.*** Schematic of the steps used to process, integrate, and evaluate paired single-cell GEX, ADT, and BCR data. Processed Cell Ranger outputs are downloaded directly, then GEX and ADT data are read into Seurat and undergo quality control (minimum number of features and maximum mitochondrial percentage), the standard Seurat pipeline, doublet removal, and automated cell type annotation. In parallel, AIRR data are processed with Immcantation (via Change-O) or nf-core/airrflow, then combined into a single BCR table with additional annotations (e.g., binned SHM frequency, gene families, amino acid properties, and novel allele checks), with cells carrying only light chains or lacking a constant region call removed. BCRs are then represented either as engineered sequence features or as numerical embeddings from protein and antibody language models (AntiBERTa2, AntiBERTy, BALM-paired, ESM2, Immune2vec). After adding AIRR information to the Seurat metadata and subsetting to cells with paired BCRs, the BCR representations are integrated with GEX either by weighted nearest neighbors (WNN) or by concatenation, and the resulting spaces are evaluated using local neighborhood consistency. Gray boxes indicate parameters, with the options used in this study underlined in blue.
